# Ecological context determines whether inherited stress responses of hosts benefit or harm herbivores

**DOI:** 10.1101/2025.10.02.680082

**Authors:** Alexandra Chávez, Amanda de Santana Lopes, Martin Lohr, Martin Schäfer, Lisa-Marie Laick, Enrico Diniz Rodrigues Batista, Veit Grabe, Vivianne Hell, Ida Faust, Shuqing Xu, Martin Kaltenpoth, Meret Huber

## Abstract

Inherited stress responses are widely assumed to be adaptive for plants by preparing offspring for renewed attack. To date, it remains unclear whether and under which conditions these responses may benefit the attacker rather than the host. Studying the interaction between the aphid *Rhopalosiphum nymphaeae* and its native host, the aquatic duckweed *Spirodela polyrhiza*, we show that aphids can either benefit or suffer from inherited stress responses, depending on metabolite identity and ecological context. In multigenerational clonal plant lineages maintained indoors, ancestral aphid herbivory increased aphid reproduction and reduced plant fitness under renewed attack, indicating that the net effect of inherited stress responses can favor the herbivore. However, not all inherited responses benefited the aphid: using genetic and chemical manipulations, we found that transgenerational priming of jasmonate enhanced aphid performance, whereas transgenerational priming of tyramine reduced it. Notably, tyramine—but not jasmonates—accumulated to higher levels across generations also in large mesocosms outdoors, albeit only within and not across growing seasons. Together, our results reveal that inherited responses of hosts can favor the attackers and highlight ecological context and life-cycle transitions as key determinants of the persistence, duration, and ecological consequences of inherited stress responses.

## INTRODUCTION

Inherited stress responses are widely assumed to enhance offspring performance under recurrent biotic and abiotic stress by allowing parents to prepare offspring for similar conditions (*1*). However, inherited responses do not necessarily reflect adaptive anticipation (*2, 3*). Instead, they may arise from physiological constraints, including reduced resource provisioning (*4*). Furthermore, in biotic interactions, antagonists can manipulate host physiology, particularly phytohormones, for their own benefit (*5, 6*), raising the possibility that inherited responses may favor attackers rather than their hosts. Yet, whether inherited responses from hosts can benefit antagonists, and under which ecological conditions such effects persist, remains poorly understood. Resolving this question is important for understanding the ecological and evolutionary consequences of stress memory in natural populations and for evaluating whether inherited stress responses can be harnessed for crop improvement.

A central challenge in evaluating the adaptive value of inherited responses is their context dependence. Effects observed under controlled laboratory conditions may not translate to natural environments (*3*), where fluctuating abiotic conditions and complex biotic interactions modulate baseline physiology and can amplify, mask, or override inherited responses (*7*). In addition, the persistence of inherited responses may depend on temporal dynamics and life-cycle transitions (*8*). Seasonal changes often trigger shifts in reproductive mode or the formation of resting stages, during which environmentally induced phenotypes or epigenetic states may be reshaped (*9, 10*). Such interruptions to continuous reproduction may restrict the duration over which responses are inherited and influence ecological interactions. Thus, assessing the ecological relevance of inherited stress responses requires determining not only whether inherited responses affect fitness, but also whether these effects persist under realistic environmental conditions and across life-cycle transitions.

Assessing the adaptive value of inherited stress responses presents additional challenges. To distinguish inherited responses from underlying genetic variation, selection, and early-life exposure effects, genetically stable, single-descendant lineages must be tracked across several generations (*11–13*). Moreover, the fitness consequences of inherited responses should be evaluated within the specific study system, since many traits, including defense-associated metabolites, can have context-dependent effects (*14*). Even canonical defense hormones such as jasmonates can enhance herbivore reproduction under certain conditions (*3*), underscoring that fitness consequences must be empirically resolved for each interaction.

A suitable system to test whether and when inherited stress responses can benefit herbivores rather than their hosts is the interaction between the waterlily aphid (*Rhopalosiphum nymphaeae)* and its native host, the giant duckweed (*Spirodela polyrhiza)*. *S. polyrhiza* is genetically stable in multigenerational experiments due to its low mutation rate and clonal reproduction (*3, 15*), in which flat, thallus-like shoots (“fronds”) are produced approximately every two days (*16*). To overwinter, *S. polyrhiza* produces vegetative resting bodies known as turions (*17*). In early summer, *S. polyrhiza* is attacked by *R. nymphaeae*, which reproduces parthenogenetically throughout the growing season (*18*).

Using this experimentally tractable system, we show that inherited host responses can either benefit or harm herbivores, depending on metabolite identity and ecological context. This study thereby challenges the view that inherited stress responses are inherently adaptive for plants and identifies environmental context and life-cycle transitions as key determinants of the persistence and fitness consequences of inherited responses.

## RESULTS

### Indoors, ancestral herbivory decreases plant fitness and benefits aphids

To test whether ancestral aphid herbivory alters plant and aphid fitness, we pre-treated eight *S. polyrhiza* genotypes for five generations with or without the aphid *R. nymphaeae*, followed by five generations of recovery. We then quantified plant and aphid fitness under control conditions and recurrent herbivory (Fig. **1A**).

**Fig. 1.**
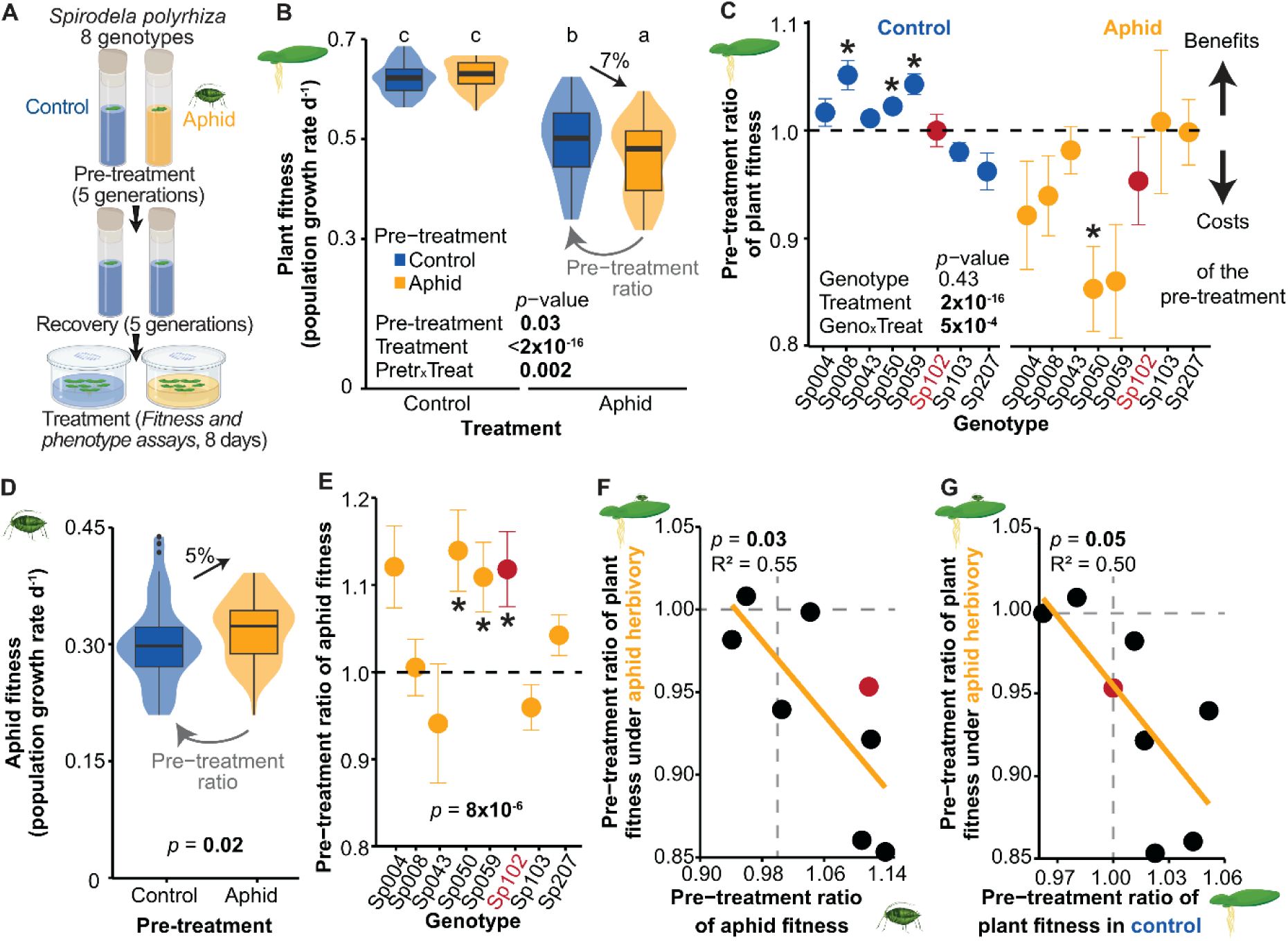
Indoors, ancestral herbivory by the waterlily aphid *Rhopalosiphum nymphaeae* boosts aphid reproduction and decreases the fitness of the duckweed *Spirodela polyrhiza*. (**A**) Schematic of the indoor transgenerational experiment considering eight *S. polyrhiza* genotypes with and without recurring aphid herbivory. N = 8 per genotype and treatment. (**B**) Across all *S. polyrhiza* genotypes, ancestral herbivory reduced plant fitness (daily population growth rates) under recurrent stress. *P*-values refer to a mixed-effects model, and letters on top indicate significant grouping by the least squares means post-hoc test. N = 58-64 (N ∼ 8 per genotype). (**C**) Pre-treatment ratios of plant fitness (fitness of aphid pre-treated plants relative to the mean fitness of control pre-treated plants) varied among genotypes. Error bars denote ± standard errors. Dashed line marks neutrality (pre-treatment ratio equal to one); asterisks denote significant deviation from neutrality (Wilcoxon rank-sum test, *P* < 0.05); *p*-value refers to a mixed-effects model. N = 5-8. (**D**) Across all *S. polyrhiza* genotypes, ancestral aphid herbivory boosted aphid fitness (daily population growth rates). *P*-values refer to the mixed-effects model. N = 58. (**E**) The aphid *R. nymphaeae* genotype Rn001 benefitted from ancestral aphid herbivory only in a subset of *S. polyrhiza* genotypes. Dashed line marks neutrality; error bars denote ± standard errors; asterisks denote significant deviations from neutrality (Wilcoxon rank-sum tests) *p*-value refers to a mixed-effects model. N = 5-8. (**F**) The more aphids benefitted from ancestral stress (increased aphid fitness pre-treatment ratios), the more the plants suffered (decreased plant fitness pre-treatment ratios). Each dot represents a genotype mean; dashed lines mark neutrality; *p*-value refers to a linear regression. N = 5-8. (**G**) The more a genotype suffered from ancestral herbivory under recurring herbivory (decreased fitness pre-treatment ratios), the better it performed when the aphids did not recur. Each dot represents a genotype mean; red dots show mean values of genotype Sp102; bold values highlight significant *p*-values; dashed lines mark neutrality; *p*-value refers to a linear regression. N = 5-8. Geno = genotype, Pretr = pre-treatment, Treat = treatment.

Unexpectedly, under recurrent stress, ancestral herbivory reduced plant fitness and increased aphid reproduction. Ancestral herbivory reduced plant fitness under recurring conditions on average by 7% (Fig. **1B**), with some genotypes suffering over 15% fitness loss (Fig. **1C**). Likewise, ancestral herbivory amplified stress-induced morphological changes (Fig. **S1A**). Notably, ancestral herbivory increased aphid reproduction on average by 5% (Fig. **1D**), with aphids reproducing up to 10% faster on certain plant genotypes (Fig. **1E**). The more a genotype suffered from ancestral herbivory under recurring stress, the more the aphids benefitted from the previous infestation (Fig. **1F**), and the more the plant benefitted when herbivory did not recur (Fig. **1G**). Together, these results demonstrate that inherited stress responses entail context-dependent fitness costs and benefits and, counterintuitively, can be maladaptive for plants under recurring stress.

### Tyramine and jasmonates are transgenerationally primed

To identify metabolites underlying inherited fitness effects, we profiled free amino acids, amines, phytohormones, and phenylpropanoids at the end of the “fitness and phenotype assays” in *S. polyrhiza* Sp102, a genotype that supported higher aphid fitness when pre-treated with the herbivore (Fig. **1E**, Fig. **S1B**). Under control conditions, ancestral herbivory elevated four metabolites, including tyramine (Fig. **2A**, left panel). Under aphid herbivory, 14 out of 46 metabolites accumulated to up to three-fold higher levels in aphid pre-treated than control pre-treated plants, most prominently tyramine and jasmonic acid (Fig. **2B**). No metabolite was suppressed by herbivory pre-treatment (Fig. **2A**, right panel).

**Fig. 2.**
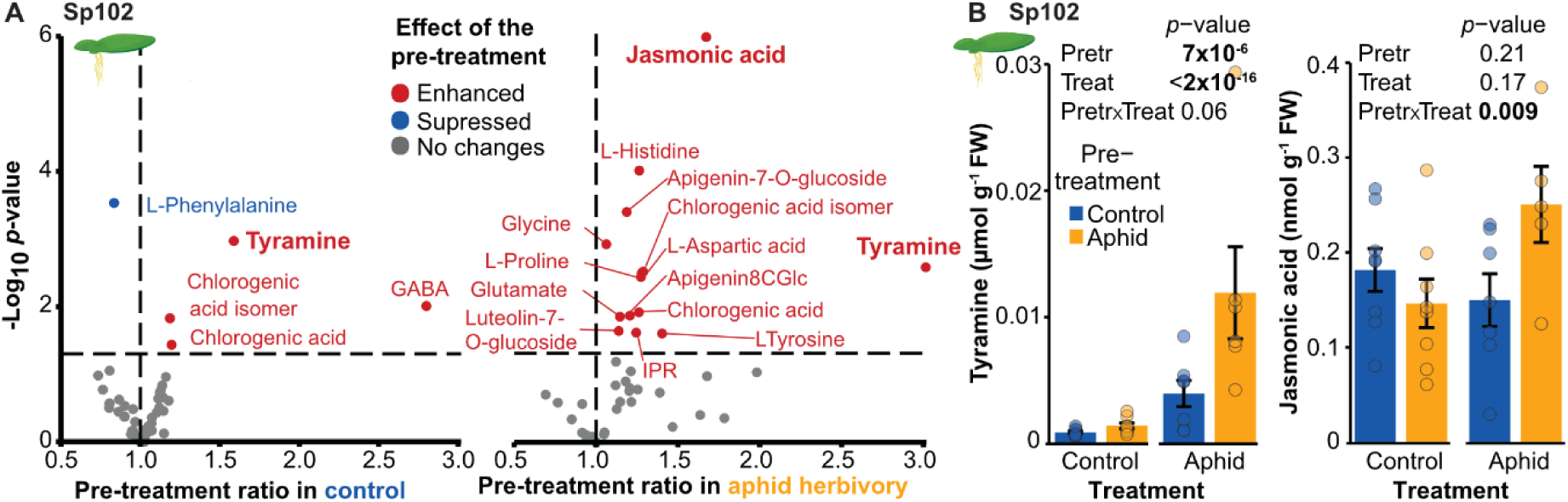
Ancestral aphid herbivory transgenerationally primes *Spirodela polyrhiza* for elevated accumulation of tyramine and jasmonate. (**A**) Volcano plot depicting metabolites that are transgenerationally plastic in the absence (left panel) and presence (right panel) of recurring aphid herbivory. Vertical dotted lines mark the neutral fold change (no pre-treatment effect), and horizontal dotted lines indicate the significance threshold (*P* < 0.05, mixed-effects model). Each point represents the mean metabolite level measured at the end of the indoor fitness and phenotype assay in genotype Sp102. N = 5-8. (**B**) Bar plot depicting the levels of tyramine (left panel) and jasmonic acid (right panel) in the presence and absence of recurring stress in genotype Sp102, measured at the end of the indoor fitness and phenotype assay. Error bars denote ± standard errors. *P*-values refer to mixed-effects models. N = 5-8. Bold values highlight significant *p*-values. GABA = γ-aminobutyric acid, iPR = isopentenyladenine riboside. FW = fresh weight, Pretr = pre-treatment, Treat = treatment.

Tyramine levels were both transgenerationally retained and primed. First-time herbivory enhanced tyramine by six-folds (Fig. **S1C**), and elevated levels were maintained across generations in the absence of stress, with herbivory pre-treated plants accumulating 60% more tyramine than control pre-treated plants (Fig. **2B**, Fig. **S1C**). Under recurring herbivory, tyramine levels tripled in plants with prior herbivory exposure in comparison to plants under first time herbivory (Fig. **2B**). Transgenerational plasticity in tyramine was not only observed in Sp102, but also in half of the other genotypes (Fig. **S1D**).

In contrast, jasmonic acid was transgenerationally primed but not retained: jasmonic acid was not induced upon first time herbivory exposure (Fig. **S1E**), and consequently, was not retained (Fig. **2B**). However, under recurrent herbivory, aphid pre-treated plants accumulated 70% higher levels of jasmonic acid compared to control pre-treated plants (Fig. **2B**). Similar patterns and magnitudes were observed for jasmonic acid-isoleucine (Fig. **S1F**).

### Transcription priming of the biosynthetic machinery may prime tyramine

To determine whether transgenerational priming of tyramine reflects regulation of its biosynthetic machinery, we identified tyrosine decarboxylases (*TYDCs*)—encoding the enzymes that convert tyrosine into tyramine (Fig. **3A**)—and quantified their expression. Based on genome annotation, we identified three putative *TYDCs* in the *S. polyrhiza* genome (Fig. **3B**, Fig. **S2A** and Table **S1**). To validate their function, we transiently expressed the genes in *S. polyrhiza* calli, using the rice *TYDC* as a positive control and the annotated *S. polyrhiza* tryptophan decarboxylase (*TDC*) as a negative control (Table **S1**). SpTYDC1 and SpTYDC2, but not SpTYDC3, catalyzed the decarboxylation of tyrosine (Fig. **3C**, Fig. **S2B**).

**Fig. 3.**
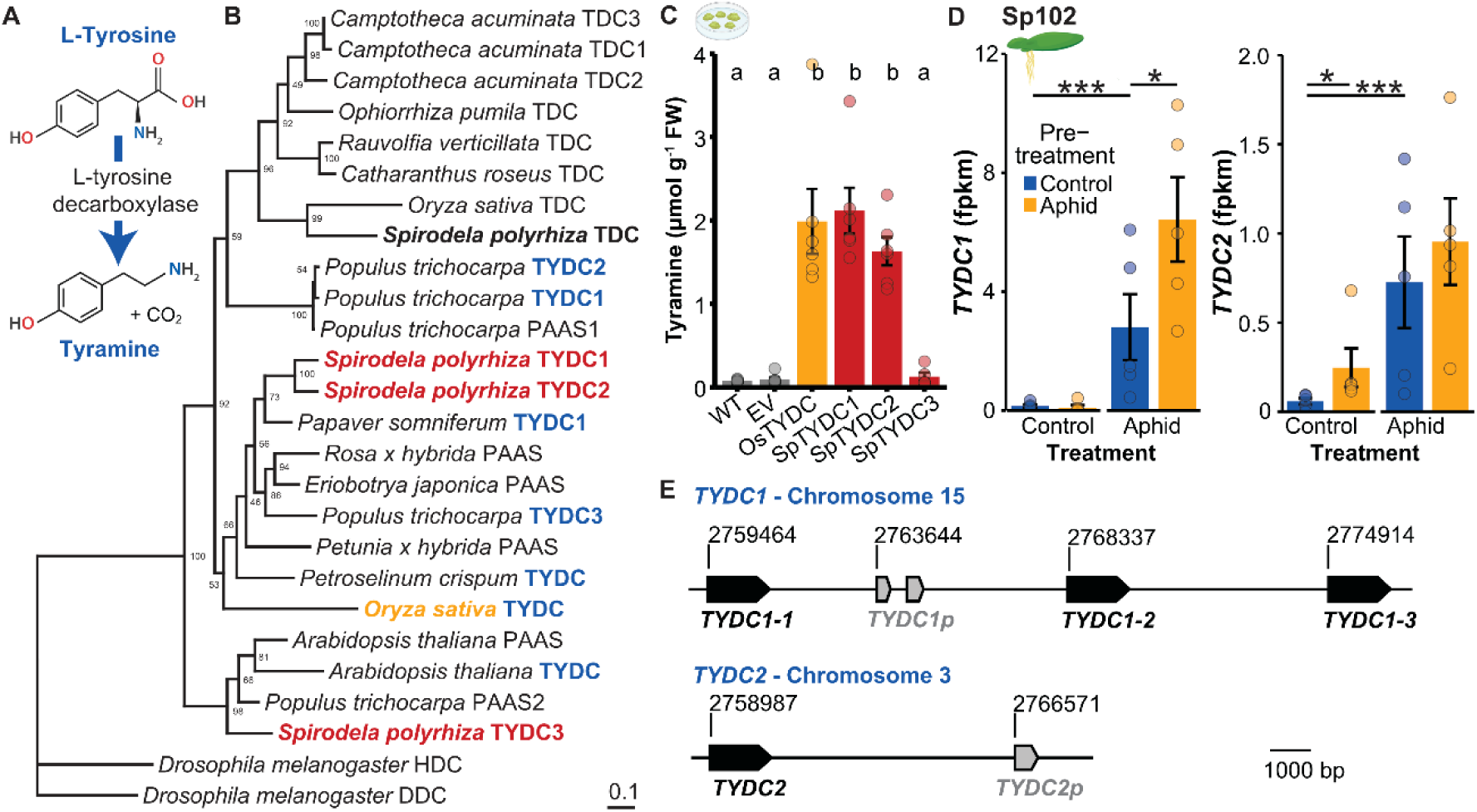
Ancestral aphid herbivory primes *Spirodela polyrhiza* to upregulate the expression of its tandemly duplicated L-tyrosine decarboxylase genes under aphid attack. (**A**) L-tyrosine decarboxylases (TYDC) catalyse the decarboxylation of L-tyrosine to tyramine. (**B**) Phylogenetic tree of characterized aromatic amino acid decarboxylases (Table S3) and their homologs in *S. polyrhiza*. L-tyrosine decarboxylases are highlighted in blue; the three *S. polyrhiza* L-tyrosine decarboxylases are shown in bold red, and the *S. polyrhiza* L-tryptophan decarboxylase in bold black. Numbers at branching points refer to bootstrap values. (**C**) Transient expression of candidate genes in *S. polyrhiza* calli revealed that TYDC1 and TYDC2 but not TYDC3 are tyrosine decarboxylases. The rice tyrosine decarboxylase (OsTYDC, (*50*)) served as a positive control. N = 6. (**D**) Ancestral aphid herbivory elevated expression of the *S. polyrhiza* L-tyrosine decarboxylases *TYDC1* and *TYDC2*, both of which are induced by aphid feeding. Asterisks denote significant differences using generalized linear models on transcriptome data derived from Sp102 plants harvested at the end of the indoor fitness and phenotype assay. N = 5. (**E**) Genomic organization and gene models of the two *S. polyrhiza TYDC* homologs. Black boxes represent predicted coding exons and grey boxes denote pseudogenes. Chromosomal coordinates (bp) are indicated above each locus. DDC = L-dopa decarboxylase, HDC = histidine decarboxylase, PAAS = phenylacetaldehyde synthase, TDC = tryptophan decarboxylase, TYDC = tyrosine decarboxylase; EV = empty vector, WT = wild type (regenerated Sp162), Os = *Oryza sativa*, Sp = *Spirodela polyrhiza*; FPKM = fragments per kilobase of transcript per million mapped reads, FW = fresh weight. * < 0.05; *** < 0.001.

Transcriptomes from the fitness assay revealed that the expression levels of *SpTYDC1* and *SpTYDC2* mirrored tyramine accumulation: both genes were more strongly expressed under aphid herbivory than under control conditions, and their expression was transgenerationally retained and primed (Fig. **3D**). Because *SpTYDC1* showed higher expression under aphid herbivory and a stronger transcriptional response to aphid pre-treatment than *SpTYDC2* (Fig. **3D**), transgenerational plasticity of tyramine is likely driven primarily by plasticity in *SpTYDC1* transcription.

Although *TYDC1* and *TYDC2* are co-regulated and share 82% amino acid similarity, the two genes are located on different chromosomes. Notably, the chromosomal segment containing *TYDC1* harbors three tandem copies of the gene along with one *TYDC1* pseudogene (*TYDC1-1*, *TYDC1-2*, *TYDC1-3*; Fig. **3E**, Table **S1**). Similarly, *TYDC2* is accompanied by a *TYDC2* pseudogene (Fig **3E**, Table **S1**). The presence of multiple functional TYDC gene copies supports the notion that tyramine plays an important function in *S. polyrhiza*.

### Tyramine suppresses aphid fitness

Tyramine may act as a plant defense through toxicity or interference with herbivore hormonal signaling but may also serve as an insect resource by contributing to metabolic pathways underlying cuticle sclerotization and melanization (Fig. **S2C**). We therefore chemically supplemented SP102 with tyramine and measured plant fitness, as well as aphid fitness, *Buchnera* titers–aphid endosymbionts that contribute to tyrosine supply(*19*)–and aphid cuticle sclerotization and melanization. Tyramine supplementation increased *in planta* tyramine levels comparable to those observed in our transgenerational experiment (Fig. **2B**, Fig. **S3A**) and alongside increased the levels of the tyrosine-derived metabolites dopamine (Fig. **S3B**). Tyramine supplementation improved plant resistance: while tyramine application reduced plant fitness under control conditions, this effect tended to be reversed under herbivory (Fig. **4A**, left panel). Furthermore, tyramine decreased aphid fitness at concentrations similar to those observed in transgenerational experiments, with dose-dependent reductions detectable after two days (Fig. **4A**, middle-left panel) and eight days of feeding (Fig. **S3C**).

**Fig. 4.**
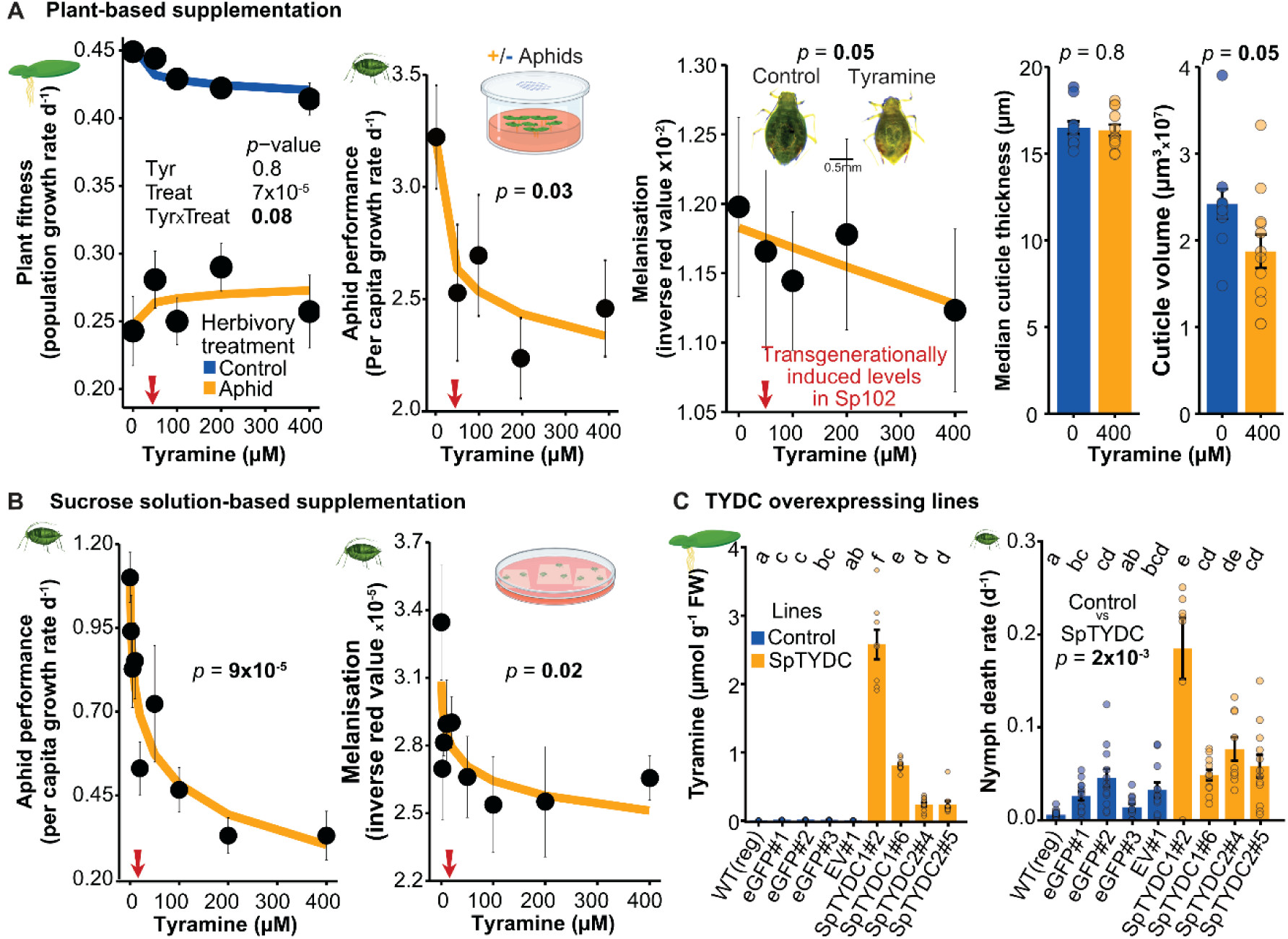
Tyramine protects *Spirodela polyrhiza* against the aphid *Rhopalosiphum nymphaeae*. (**A**) Plant fitness and aphid performance, melanization and cuticle thickness in tyramine-supplemented Sp102 plants. N = 11-12 per concentration. Left panel: Increasing tyramine reduced plant fitness (daily population growth rates over eight days) under control conditions but tended to increase it under aphid herbivory. *P*-values refer to a mixed-effects model. Middle-left panel: Aphid performance (per capita reproduction after two days of feeding) declined with increasing levels of tyramine. *P*-value refers to a linear model on log-transformed data. Middle-right panel: Tyramine supplementation decreased cuticle melanization (inverse red color intensity) in aphids after eight days of feeding. *P*-value refers to a linear model. Right panel: Tyramine supplementation did not alter aphid cuticle thickness but decreased cuticle volume. *P*-values refer to Student’s *t*-test. (**B**) Aphid fitness (per capita growth rate, left panel) and cuticle melanization (right panel) of aphids feeding for five consecutive days on 35% sucrose solutions containing 0-400 µM tyramine. N = 5 per concentration. *P*-values refer to a linear model on log-transformed data. For (**A-B**): the red arrow denotes the *in planta* tyramine concentration equivalent to that measured in Sp102 after recurring indoor aphid herbivory. Dots represent mean values per concentration, and error bars denote ± standard errors. (**C**) Tyramine levels and nymph death rate on transgenic *S. polyrhiza* plants that stably overexpress *SpTYDC1* and *SpTYDC2*. Left panel: tyramine accumulation in shoots of TYDC transgenic lines and controls (regenerated wild type, eGFP, empty vector). Right panel: death rate of nymphs over seven days of assays. Independent transgenic lines are indicated by hash numbers; error bars denote ± standard errors; *p-*values refer to mixed-effects models, and letters on top indicate significant grouping by the least squares means post-hoc test. N = 12 per line. Bold values highlight significant *p*-values. TA = tyramine treatment, Treat = herbivory treatment, WT = wild type, eGFP = enhanced green fluoresce protein, EV = empty vector, TYDC = tyrosine decarboxylase, Sp = *Spirodela polyrhiza*.

In contrast, tyramine was unlikely to serve as a nutritional resource for aphids: tyramine supplementation did not alter *Buchnera* load (Fig. **S3D**). Furthermore, tyramine supplementation reduced cuticle melanization of aphids, with aphids appearing redder rather than darker (Fig. **4A**, middle-right panel). In line, tyramine supplementation did not alter cuticle thickness or cuticle-to-body volume ratio (Fig. **4A**, right panel; Fig. **S3E**). Only cuticle volume was slightly reduced (Fig. **4A**, right panel), likely reflecting smaller body size (Fig. **S3E**).

To test whether tyramine is sufficient to reduce aphid fitness and cuticle melanization, we fed aphids with a sucrose solution supplemented with tyramine (Fig. **4B**). As expected, tyramine reduced aphid performance and cuticle melanization across the five days of feeding: already 2 µM of tyramine—4% of *in planta* tyramine levels (Fig. **S3A**)—reduced aphid reproduction and melanization (Fig. **4B**).

To assess whether endogenous production of tyramine alters plant and aphid fitness, we subjected aphids to transgenic *S. polyrhiza* lines that constitutively overexpressed either *SpTYDC1* or *SpTYDC2*, which increased basal tyramine levels up to 700-folds compared to transgenic controls (Fig. **4C**, left panel). While overexpressing *SpTYDC1* and *SpTYDC2* did not improve plant fitness under aphid herbivory—likely due to inherently lower population growth rates of the TYDC-overexpressing lines (Fig. **S4A**)—overexpression doubled mortality of nymph and adult aphids (Fig. **4C**, right panel; Fig. **S4B**), and decreased aphid fitness on TYDC plants compared to transgenic controls (Fig. **S4C**). Repeating the experiment showed similar results (Fig. **S4D**). Taken together, these data support the notion that tyramine protects plants against its aphid herbivore.

### Jasmonate priming benefits aphid fitness

Since transgenerational plasticity in tyramine cannot explain the increased aphid fitness upon recurring herbivory, we studied another transgenerationally plastic metabolite: jasmonic acid. To assess whether jasmonates alter plant and aphid fitness, we externally applied methyl-jasmonate for three days to Sp102, and subsequently measured aphid performance in the absence of methyl jasmonate. Surprisingly, performance of aphids increased at mild jasmonate doses (Fig. **5A**, left panel). The increase in aphid reproduction may be explained by a 30% rise in free amino acid pools (Fig. **5A**, right panel). We repeated the experiment including higher methyl-jasmonate levels, which showed that mild but not high jasmonate doses increase aphid performance (Fig. **S5A**). To corroborate these results, we continuously applied methyl jasmonate to plants and measured aphid performance. Accordingly, mild methyl-jasmonate doses increased aphid performance, whereas high doses reduced it (Fig. **5B**, Fig. **S5B**). Consequently, high jasmonate doses benefitted plant fitness under aphid herbivory (Fig. **S5B**). We repeated the continuous supplementation with two jasmonate doses, corroborating that mild doses increased aphid fitness (Fig. **S5C**). Taken together, these data indicate that mild but not high levels of methyl-jasmonate benefit aphid reproduction.

**Fig. 5.**
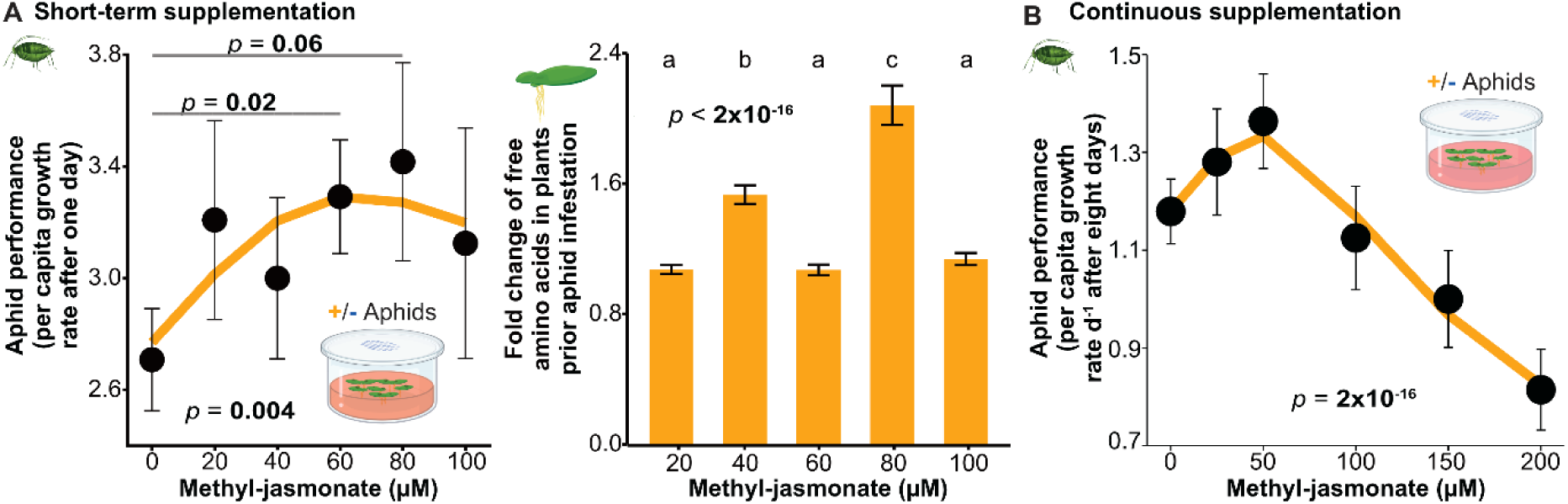
Mild methyl-jasmonate induction enhances aphid performance and increases free amino acid levels in *Spirodela polyrhiza*. (**A**) Short-term methyl-jasmonate supplementation. Left panel: Supplementing mild doses (60–80 µM) of methyl-jasmonate to *S. polyrhiza* genotype Sp102 increased the performance of the aphid *R. nymphaeae* genotype Rn001. Aphid performance (per capita growth rates) was measured over 24 h in the absence of methyl-jasmonate following three days of methyl-jasmonate induction on *S. polyrhiza*. Dots represent mean values and error bars denote ± standard errors; horizontal bars indicate significant pairwise comparisons and *p*-values above the bars refer to mixed-effects models; *p*-value at the bottom refers to a mixed-effects model with natural spline. N = 8 per concentration. Right panel: Free amino acid levels increased at moderate methyl-jasmonate doses. Amino acids were quantified after three days of methyl-jasmonate induction and before aphid infestation. Fold changes were calculated relative to solvent controls (0 µM methyl-jasmonate). N = 6–8 per concentration and amino acid. *P*-value refers to a mixed-effects model, and letters on top were obtained by the least squares means post-hoc test. (**B**) Continuous methyl-jasmonate induction. Methyl-jasmonate concentrations above 100 µM decreased aphid performance (per capita growth rates) on *S. polyrhiza* Sp102 plants. Performance was measured by counting adult aphids after eight days of free reproduction on plants exposed to 11 days of continuous methyl-jasmonate induction. Dots represent mean values and error bars denote ± standard errors; *p*-value refers to a mixed-effects model with natural spline. N = 16 per concentration. Bold values highlight *p*-values. MeJA = methyl-jasmonate, Treat = herbivory treatment.

### Plants overcome maladaptive plasticity in natural conditions

To determine whether transgenerational plasticity observed under controlled conditions persists under natural conditions, we performed reciprocal, transgenerational transplant experiments in outdoor mesocosms in which large Sp102 populations experienced contrasting levels of aphid herbivory over two consecutive growing seasons (Fig. **6A**, Fig. **S6A**). Within the first season, tyramine accumulated to higher concentrations in herbivore-exposed populations and their overwintering turions (Fig. **S6B**). In contrast, jasmonate levels were unaffected by herbivory in plants and turions throughout the season (Fig. **S6C**).

**Fig. 6.**
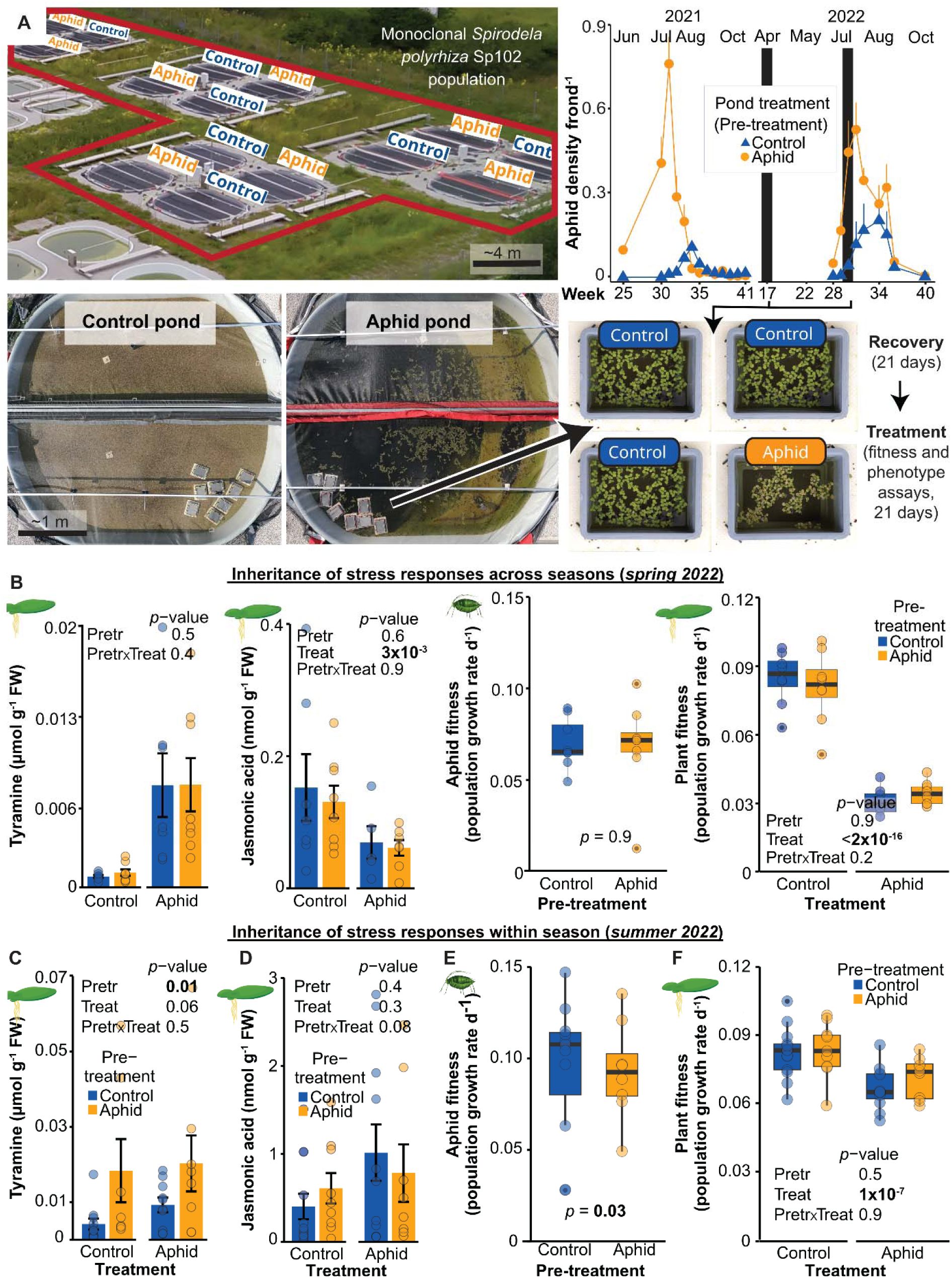
Outdoors, transgenerational tyramine priming persisted, whereas transgenerational jasmonate priming and its costs disappeared. **(A)** Experimental setup of outdoors mesocosms in which monoclonal populations of *S. polyrhiza* genotype Sp102 were grown for two consecutive growing seasons with contrasting levels of the aphid *R. nymphaeae* genotype Rn001. Upper-left panel: Experimental ponds are outlined in red. N = 8 mesocosms per treatment. Upper-right panel: The time series (from (*27*)) shows aphid population densities across the two growing seasons. Vertical black bars indicate the calendar weeks when plants were transferred to an aphid-free recovery phase. Bottom-left panel: Representative aerial views of a control and an aphid pond, each containing floating boxes used for reciprocal transplant experiments. Bottom-right panel: Schematic of the reciprocal transplant experiment, illustrated by opened floating boxes in which assays were conducted. In spring and summer 2022, plants were transferred into floating boxes and grown for three weeks under aphid-free conditions (recovery phase). Offspring produced during this period were then subjected to a three-week fitness and phenotype assay in the presence and absence of aphids. (**B**) Transgenerational plasticity in metabolite concentrations and fitness of host and herbivore do not persist across seasons. Across-season transgenerational plasticity was measured in spring 2022. Error bars refer to ± standard errors; *p*-values refer to mixed-effects models. N = 8-13 samples per treatment. (**C-F**) Within-season transgenerational plasticity increases tyramine concentrations, consistently decreasing aphid fitness. Within-season transgenerational plasticity was measured in summer 2022. Error bars refer to ± standard errors; *p*-values refer to mixed-effects models. N = 8-13 samples per treatment. Bold values highlight significant *p*-values. Pretr = pre-treatment, Treat = treatment. FW = fresh weight.

However, these aphid-induced metabolite levels did not persist across seasons. Spring-collected turions did not differ in tyramine and jasmonates accumulation, except that jasmonic acid-isoleucine was increased by 44% following ancestral aphid herbivory (Fig. **S6D**). Furthermore, plants germinated from turions did not differ in tyramine or jasmonate concentrations, plant fitness, morphology, or aphid fitness, regardless of whether aphids were re-introduced (Fig. **6B**, Fig. **S6E-F**), indicating that overwintering resets herbivore-induced metabolic states.

Within a growing season, transgenerational effects persisted. In the second season, ancestral herbivory increased tyramine levels both in the presence and absence of recurring herbivory (Fig. **6C**). Similarly, ancestral herbivory reduced frond size (Fig. **S6G**). In contrast, ancestral herbivory did not affect jasmonates in any environment (Fig. **6D**, Fig. **S6H**). In line, aphids no longer benefitted from ancestral herbivory and instead produced fewer offspring on ancestrally exposed than on non-exposed plants (Fig. **6E**). Consistently, ancestral herbivory did not reduce plant fitness under recurring herbivory (Fig. **6F**).

### Inherited stress responses are no longer detectable after turion formation

Because transgenerational plasticity in tyramine disappeared across growing seasons, we tested whether turion formation resets inherited responses. We germinated Sp102 turions that were collected during the indoor transgenerational experiment after three generations of recovery and quantified tyramine levels in the resulting lineages. Although tyramine levels showed similar directional patterns before and after turion formation, differences between control and aphid pre-treated plants were no longer detectable (Fig. **S6I**). Together, these results indicate that in natural environments stress responses are inherited within seasons but not across years because of the formation of resting bodies.

## DISCUSSION

Inherited stress responses is often interpreted as an adaptive mechanism preparing descendants for recurring stress (*1*). Our findings challenge this view by showing that these responses can be either adaptive or maladaptive for the plant, depending on metabolite identity and environmental context. Thus, inherited stress responses is not inherently adaptive under recurrent stress but may instead reflect physiological processes whose fitness consequences depend on ecological context.

One mechanism generating non-adaptive outcomes is the transgenerational priming of hormonal signaling that antagonists can exploit. In our study, aphids benefitted from ancestral infestation, possibly because mildly elevated jasmonates increased the availability of free amino acids, which are often limiting for aphids (*19*). Jasmonates may, in principle, be actively manipulated by aphids—for instance through the vertical transfer of effector proteins or non-coding RNAs (*20, 21*). Yet, since jasmonates were transgenerationally primed even when the initial stress was copper excess (*3*), inherited jasmonate levels likely reflect a stress-induced physiological response from which herbivores benefit, rather than active manipulation of the host.

Notably, mild jasmonate priming and its associated fitness costs for plants disappeared outdoors, whereas transgenerational increases in tyramine, which was defensive, persisted. Jasmonate priming may be lost outdoors because jasmonates are not consistently induced by aphids. In addition, plants accumulated fivefold higher jasmonate levels outdoors than indoors. Under such elevated baseline levels, subtle transgenerational increases are likely masked or become biologically irrelevant. In contrast, tyramine was strongly and consistently induced by aphids and showed pronounced transgenerational increases of up to three-folds following ancestral aphid exposure. Large and consistent responses likely exceed background physiological variation, allowing transgenerational effects to persist across environments and reduce aphid performance. Although tyramine has rarely been implicated as a defense metabolite against herbivores (*22*), our data shows that tyramine – and its transgenerational response - constitutes an underappreciated plant defense mechanism. Together, these findings suggest that outdoor conditions do not uniformly erase stress memory but filter transgenerational responses according to their induction strength and relative to their background physiological variation.

Outdoor conditions may not only limit the persistence of transgenerational effects by shifting baseline physiology but also by inducing life-cycle transitions (*1, 8*). In this study, transgenerational increases in tyramine disappeared across seasons and were no longer detectable in indoor lineages after regeneration from turions, indicating that life-cycle transitions reset inherited physiological states. The loss of inheritance is consistent with extensive epigenetic reprogramming during turion formation (*10*). Although epigenetic reprogramming in turions, as also observed in other dormancy bodies including seeds (*9*), is not specific to prior stress exposure, such genome-wide reorganization likely resets previously acquired epigenetic marks. Because life-cycle transitions are commonly triggered by seasonal change and clonal reproduction is widespread among flowering plants (*23*), transgenerational plasticity is likely to be ecologically relevant within growing seasons; yet its long-term ecological and evolutionary consequences may be constrained by recurrent life-cycle transitions.

Together, our results show that transgenerational plasticity in plant metabolism can have opposing ecological consequences, with some metabolites promoting aphids at the expense of the host, whereas others suppress herbivores while protecting the plant. Whether these transgenerational effects manifest in nature likely depends on whether environmental variation masks inherited signals and triggers life-cycle transitions. Transgenerational plasticity should therefore be viewed less as a reliable mechanism of adaptive anticipation and more as stress-induced physiological response whose ecological relevance depends on environmental context, signal strength, and continuous reproduction.

## MATERIALS AND METHODS

### Plant and aphid growth conditions

For the indoor experiments, we grew plants and aphids inside growth cabinets (L4Z-TDL+, poly klima GmbH, Freising, Germany) operating at 28 °C, 150 µmol photons m^-2^ s^-1^ or at 26 °C, 135 µmol photons m^-2^ s^-1^ for transgenic plants, with a 16:8 h light/dark cycle. Plants were cultivated in N-medium (*24*), except experiments involving transgenic plants, for which we used 0.5 Schenk and Hildebrant (SH) medium (Duchefa Biochemie, Haarlem, The Netherlands) (*25*). To minimize microbial contamination in non-transgenic plants, we surfaced-sterilized them at the beginning of experiments, after which the plants were grown under axenic conditions for four weeks in 250 mL Erlenmeyer flasks (Fisher Scientific GmbH, Schwerte, Germany) before starting the experiments. Transgenic plants were kept sterile during pre-cultivation within MagentaTM GA-7 boxes (no.: V8505; Sigma-Aldrich, St. Louis, USA). Aphid nymphs of the strain Rn001 (*26*) were cleared of surface fungi and algae by rearing them on sterile plants for at least five consecutive generations; during this time, the insects were sprayed every week with a copper-based antifungal solution commonly used in aquaria (Sera mycopur, Heinsberg, Germany). The resulting aphids were then acclimated for one month on Sp050 or Sp102 sterile plants for the transgenerational or the induction assays, respectively, in autoclaved 2.6 L boxes (Lock & Lock, Seul, South Korea) filled with 1 L N-medium. The boxes were covered with lids in which an area of 25 cm^2^ was excised and covered with a metal mesh (meshopening 0.3 mm, wire diameter 0.2 mm; Haver & Boeker, Oelde, Germany) for gas exchange.

For the outdoor experiment, we used the outdoor mesocosms described in Schäfer et al(*27*). To carry out transgenerational transplant experiments, we used the strain Rn001. We reared the aphids in 2.6 L boxes (Lock & Lock) filled with 1 L N-medium within walking chamber maintaining 26 °C, 135 µmol photons m^-2^ s^-1^ and a 16:8 h light/dark cycle.

### Statistical analysis

All data were analyzed in R v4.4.0 (*28*). As *fitness* metric in assays where at least two new generations were produced (F1 and F2), we calculated relative growth rates (*29*), hereafter “population growth rates”, based on the increase in plant surface area and aphid number, shown in equation (1). As *performance* metric in assays where only one new generation was produced (F1)—particularly applied for aphids, we calculated the daily per capita reproduction rate, defined as the number of nymphs produced per parent per day(*30*), equation (**2**). When using transgenic plants, we estimated the mortality incidence density, hereafter “*death rates*”, defined as the number of dead aphids divided by the total aphid density–time at risk(*31*). Density–time at risk was estimated as half of the dead and alive produced aphids during the assays multiplied by the duration of the assay (in days); equation (**3**). This metric standardizes mortality by aphid density and time at risk, providing a per capita death rate that accounts for changes in population size during the assay. To assess plant morphology, we measured surface area per frond (mm^2^), fresh weight per frond (mg) and surface area per fresh weight (mm^2^ mg^-1^). Additionally, we calculated the *pre-treatment ratios* by dividing each aphid pre-treated plant measurement by the mean value of the control pre-treated plants within each treatment environment(*32*), equation (**4**).

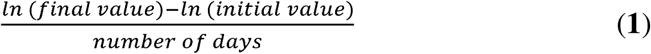

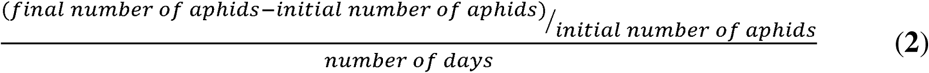

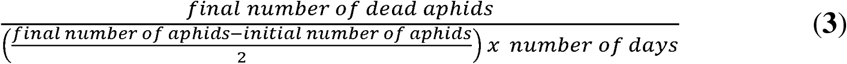

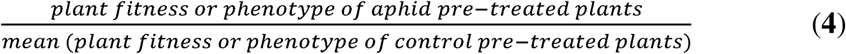

Mixed-effect models were performed with the package glmmTMB v1.1.9 (*33*). In the absence of random factors, we performed simple linear models and generalized models with lm and glm, respectively (*28*). *P*-values of the fixed-effects models were estimated with analysis of variance (Anova), using Wald chi-square of the package car (*28*). R^2^ values were obtained with ggpmisc v0.5.6 (*34*). Post-hoc *P*-values were calculated with emmeans v1.10.2(*35*) and letters were obtained with multcomp v1.4.25 (*36*). To assess the fitted models, we used the package DHARMa v0.4.6 (*37*) and effects v4.2.2 (*38*). To estimate deviation from neutral, we performed Wilcoxon Rank-Sum tests of the pre-treatment ratios per environment against one (mu=1). We displayed plots with ggplot2 v3.4.3 (*39*) and organized data with readxl v1.4.2 (*40*), data.table v1.14.8 (*41*), tidytext v0.4.1 (*42*) and dplyr v1.1.2 (*43*).

### Metabolite quantification

Amino acids, amines, phenylpropanoids and phytohormones were analysed via LC-MS as described previously (*3, 44*). In short, we ground the flash-frozen plant material of each replicate to a fine powder using a MM301 Mixer Mill (Retch, Haan, Germany). Subsequently, we extracted 20 mg ground samples using acidified methanol (MeOH:water:formic acid 15:4:1□v/v/v) containing the internal standards for the quantification of phytohormones and phenylpropanoids. We then diluted an aliquot of this extract 1:100 in an aqueous solution of isotope-labelled amino acids (algal amino acid mixture-^13^C-^15^N; Sigma-Aldrich) to quantify free amino acids and amines. The remaining extract was purified and partially concentrated via two solid-phase extraction steps using Chromabond HR-X and HR-XC columns (Macherey-Nagel, Düren, Germany) to quantify phytohormones and phenylpropanoids. The MRM-settings, ESI-settings, the gradient program and column oven settings were used as described previously (*3, 44*), with the addition of kynurenine, kynurenic acid, serotonin, L-DOPA, dopamine, GABA and melatonin to method 1A (Table **S2**). Analytes were quantified based on internal standards. The peaks from all metabolites were integrated with LabSolution Insight Version 4.0 SP6 (Shimadzu).

### Transgenerational experiment with eight *S. polyrhiza* genotypes

#### Pre-treatment, recovery, and fitness and phenotype assays

To assess whether ancestral aphid herbivory alters duckweed fitness, phenotype, and aphid reproduction, we exposed single-descendant lineages of *S. polyrhiza* to five generations of *R. nymphaeae* herbivory or control conditions (“pre-treatment”), followed by five generations under control conditions (“recovery”). Subsequently, we measured plant and aphid fitness, as well as plant morphology, both in the presence and absence of herbivory across eight days of free growth (“treatment”) (Fig **1A**), with eight replicates per genotype and treatment. For *S. polyrhiza*, we used eight range-wide sampled genotypes from each of the four genetic clusters (*15, 45*)–Sp004 (clone 7498) and Sp008 (clone 8683) from America, Sp043 (clone 9497) and Sp059 (clone 9509) from India, Sp050 (clone 0090) and Sp207 from Southeast Asia, and Sp102 and Sp103 from Europe. For *R. nymphaeae*, we used the German Rn001 strain (*26*).

Prior to the experiment, we acclimated the plants by propagating individual fronds as single-descendant lineages for four generations. We then began the pre-treatment phase by allocating the first and second offspring of each plant equally into a control and a herbivory treatment, the latter consisting of three adult aphids per plant. Lineages were propagated for five generations under these conditions (“pre-treatment”), then for an additional five generations under control conditions (“recovery”), resulting in generation 10. Throughout the acclimation, pre-treatment and recovery phase, each plant was grown individually in a 30 mL polypropylene tube (Fisher Scientific, Waltham, USA) filled with 25 mL N-medium and closed with a sterilized foam plug (CarlRoth, Karlsruhe, Germany). At each generation, as soon as an offspring frond had fully expanded, we transferred that offspring into a fresh tube (*13*). To distinguish parent and offspring, we marked the parents with a permanent marker pen (Stabilo OHPen Universal, Heroldsberg, Germany).

To minimize positional and clustering effects within 24-well racks, we used a balanced rotational design in which four genotypes per rack were systematically permuted across left-to-right positions. Across nine unique combinations, each genotype occupied all rack positions and was distributed across multiple rack contexts, reducing spatial bias. To avoid birth-order bias, first- and second-generation offspring plants were allocated in equal proportions to pre-treatment and treatment phases. Within each replicate, birth order (first vs. second offspring) was alternated between control and aphid environments to ensure balanced representation across experimental conditions and to avoid systematic bias.

To assess plant fitness and phenotype, as well as aphid fitness, we allocated the first and second offspring of generation 10 equally to control conditions or herbivory, with three adult aphids per plant (“treatment” in the “phenotype and fitness assay”). These plants were grown in 250 mL transparent polypropylene beakers filled with 150 mL N-medium and covered with perforated transparent lids (Fig **1A**). Medium was replaced every 3-5 days to overcome any contamination caused by the aphids. After eight days, we counted the number of aphids and subsequently removed them. We then harvested the plants by briefly drying them with tissue paper, weighing them and immediately flash-freezing them in liquid nitrogen. To quantify surface area and frond number, we captured images at the beginning and end of each assay using a camera box installation with a webcam (HD Pro Webcam C270, Logitech, Lausanne, Switzerland; webcam software 2.12.8). We analyzed surface area of plants with the package pliman v2.1.0 (*46*) and counted fronds with dotdotGoose v1.5.3 (*47*). All samples were stored at −80 °C until metabolite and RNA extractions.

To assess whether the pre-treatment affects plant and aphid fitness, as well as plant morphology, across the eight genotypes, we fitted mixed-effects models with pre-treatment, treatment and their interaction as fixed effects, and replicate, genotype, birth order and the rack as random effects: *Variable ∼Pre-treatment*Treatment +(1|Replicate)+(1|Genotype)+(1|BirthOrder)+(1|Rack).* As variables, we used plant and aphid population growth rates, surface area per frond, biomass per frond, and surface area per biomass. To analyze whether the pre-treatment ratios depended on genotype and treatment, we used a mixed-effects model with treatment as a fixed effect and lineage as a random effect: *Variable ∼Genotype*Treatment +(1|Lineage)*. When analyzing aphid populations growth rates and their pre-treatment ratios, we dropped the treatment fixed effect. To identify genotypes in which the pre-treatment ratios of plant and aphid fitness differed from neutral, we used Wilcoxon Rank Sum tests.

Using genotype means of pre-treatment ratios, we fitted linear models to examine two relationships: first, whether the plant fitness pre-treatment ratios under aphid herbivory correlated with the corresponding aphid fitness pre-treatment ratios; and second, whether the plant fitness pre-treatment ratios measured under control conditions correlated with those measured under herbivory.

#### Analysis of metabolites

To determine whether the pre-treatment altered metabolites accumulation, we fitted linear mixed-effects models separately within control and aphid treatments, using the pre-treatment as a fixed effect and the lineage and genotype as random effect: *Metabolite∼Pre-treatment +(1|Lineage)+(1|Genotype)*. Similarly, to assess whether metabolites accumulation is transgenerationally primed (interaction of pre-treatment and treatment), we fitted a model with the full factorial design*: Metabolite ∼Pre-treatment *Treatment+(1|Lineage)+(1|Genotype)*. When analyzing Sp102, we dropped the genotype random effect in both mixed-effect models.

### Identification of L-tyrosine decarboxylases in *S. polyrhiza*

#### Genome mining and phylogeny of TYDC candidates

To identify the candidate genes involved in tyramine biosynthesis, we first mined the genome annotation of the *S. polyrhiza* genotype 7498 (Sp004) (*45*) for loci annotated as L-tyrosine decarboxylase (*TYDC*). This search returned three candidates: *TYDC1* (*SpGA2022_016537*), *TYDC2* (*SpGA2022_004481*), and *TYDC3* (*SpGA2022_012613*). Because tyrosine decarboxylases are sometimes mis-annotated as tryptophane decarboxylases, we also retrieved the genes annotated as tryptophane decarboxylases (*TDCs*): *SpGA2022_000740*, *SpGA2022_000739*, and *SpGA2022_000738*. To confirm that the open reading frames of these genes are complete and to identify any tandem duplicates, we searched for homologs in two additional *S. polyrhiza* genome assemblies within https://www.ncbi.nlm.nih.gov/datasets/genome/?taxon=29656-clone 7498 (*48*) and clone 9512 (*10*)–using amino-acid sequence homology (TBLASTN). Gaps in the *TYDC* region of the 7498 assembly prompted us to focus subsequent analyses on clone 9512. Finally, we aligned each predicted coding sequence to the clone 9512 assembly with ClustalW in BioEdit Sequence Alignment Editor v7.0.9.0 (*49*).

To support the functional annotation of the predicted L-tyrosine and L-tryptophan decarboxylases, we constructed a phylogenetic tree with characterized L-tyrosine and L-tryptophan decarboxylases from other species (Table **S3**). We constructed two phylogenies of nucleotide sequences with RAXML v8.2.10 for multi-threading, using advanced vector extensions; one phylogeny included the main annotated genes and the second included all the identified copies of each gene. We applied a rapid bootstrap analysis and searched for best-scoring maximum likelihood trees, using the Whelan and Goldman substitution model with gamma-distributed rate variation and 100 bootstrap replicates. Finally, to identify relatedness among the genes and gene copies, we built an identity matrix using the default similarity matrix BLOSUM62 of BioEdit.

### Transient expression of TYDC candidates in S. polyrhiza calli

To validate the role of the candidate decarboxylases, we transiently overexpressed the candidate genes in *S. polyrhiza* calli, using a well-characterized TYDC (*50*) (GenBank accession number AK065830.1) from rice as a control. To this end, we first amplified the TYDC sequences using shoots of *S. polyrhiza* genotype Sp102 cultivated for seven days in N-medium with and without aphid herbivory and leaves of *O. sativa* japonica cv. Kitaake. We snap froze the plant material in liquid nitrogen and ground it to a fine powder using a laboratory mill (Retsch, Haan, Deutschland). Then, we extracted total RNA using NucleoSpin RNA Plant (Macherey-Nagel), following manufacturer’s instruction, and synthetized the cDNA using SuperScript™ III Reverse Transcriptase (Invitrogen, Lofer, Österreich). Next, we amplified the coding sequences (CDSs) of *OsTYDC*, *SpTYDC1*—sequenced taken from the paralogue *SpTYDC1-1*, *SpTYDC2*, *SpTYDC3* and *SpTDC* using gene-specific primers containing 5′ overlap sequences for DNA assembly, and the Phusion™ High-Fidelity DNA Polymerase (Invitrogen) protocol. The overlap regions enabled ligation of PCR products into the vector backbone using the NEBuilder® HiFi DNA Assembly protocol (New England Biolabs). For *OsTYDC*, the forward primer was 5′-gttgtttggtgttacttctgcagaaaatggcaccaccctcgcactg-3′ and the reverse primer was 5′-gccatggccgcgggatatcttacttttagccacgtttacgaaggattgacg-3′. For *SpTYDC2*, the forward primer was 5′-gtttggtgttacttctgcagaaactagtcaccttaagttcaacaactcagagc-3′ and the reverse primer was 5′-tggccgcgggatatcttacttgtacacttcgacaaacagtcgaaccctctc-3′. For *SpTYDC1*, the forward primer was 5′-gtttggtgttacttctgcagaaactagtatgggcagcatccaaaccgattg-3′ and the reverse primer was 5′-tggccgcgggatatcttacttgtacattctggtctccctacgaccttga-3′. For *SpTDC*, the forward primer was 5′-gtttggtgttacttctgcagaaactagtatgggaagccttgatgccaactg-3′ and the reverse primer was 5′-tggccgcgggatatcttacttgtacacgtaatcaccgggcagcttcc-3′. For *SpTYDC3*, the forward primer was 5′-gtttggtgttacttctgcagaaactagtatggacgccgaacagctcag-3′ and the reverse primer was 5′-tggccgcgggatatcttacttgtacatcatggttcgttgctccccaa-3′.

Second, we ligated the coding-sequences obtained by RT-PCR into pZmUBQ::eGFP vector (https://www.plasmids.eu/plasmid/viewPlasmid/869 (*25*))–previously linearized by *SpeI* and *BsrGI* (New England Biolabs) digest–using NEBuilder^®^ HiFi DNA Assembly (New England Bioloabs) protocol. This procedure replaced the eGFP sequence in the expression cassette *pZmUBQ::**eGFP**::t35S* by the candidate genes. Therefore, the resulting constructs contained the candidate genes between the *Zea mays* ubiquitin promoter (pZmUBQ) and the cauliflower mosaic virus 35S terminator (t35S)–*pZmUBQ::**OsTYDC**::t35S*, *pZmUBQ::**SpTYDC1**::t35S*, *pZmUBQ::**SpTYDC2**::t35S*, *pZmUBQ::**SpTYDC3**::t35S*, and *pZmUBQ::**SpTDC**::t35S*–each combined with the hygromycin resistance cassette *pNOS::hptII::tNOS*, where hptII refers to hygromycin B phosphotransferase conferring hygromycin resistance, pNOS denotes the nopaline synthase promoter and tNOS, the nopaline synthase terminator (Fig. **S4E**). The original pZmUBQ::**eGFP**::t35S construct served as a positive control. Finally, we produced a plasmid bearing only the selectable marker (empty vector, **EV**) by removing the whole eGFP expression cassette via PCR (Phusion™ High-Fidelity DNA Polymerase; Invitrogen) and ligating the PCR product ends using NEBuilder^®^ HiFi DNA Assembly (New England Bioloabs) (Fig. **S4E**).

The plasmids generated in this study have been deposited in the European Plasmid Repository (www.plasmids.eu). The repository codes will be provided before publication.

Third, we used calli originating from Sp162 fronds and transformed these calli via *Agrobacterium tumefaciens* (*25*). In short, we first induced calli by placing *S. polyrhiza* colonies upside down onto callus induction medium (CIM)–Murashige & Skoog (MS) medium including vitamins (Duchefa Biochemie) supplemented with 1% (w/v) sucrose, 500 mg L^-1^ 2-morpholinoethanesulfonic acid monohydrate, 50 mg L^-1^ citric acid monohydrate, 100 mg L^-1^ polivinilpirrolidona average mol wt 40000, 0.2 mg L^-1^ 2,4-Dichlorophenoxyacetic acid, 0.4 mg L^-1^ thidiazuron, and 0.4% (w/v) gelrite, pH 5.8–within long day conditions (16:8h light/dark), 10 μmol photons m^−2^ s^−^1 and 26 °C. After four weeks, we transferred the calli originated from the meristematic pockets to fresh CIM plates and further propagated them for 8-12 weeks (medium refreshed every 2-3 weeks) until the calli became light-green and fast-growing.

Then, we transformed *A. tumefaciens* strain EHA105 chemically competent cells by heat shock and screened them by PCR (*51*). The cells were cultivated in liquid LB, harvested by centrifugation, resuspended in infection medium (10 mM magnesium chloride, 5% sucrose, 200 µM acetosyringone) to the OD of 0.7, and incubated at room temperature for 1 hour with mild agitation. Afterwards, we transferred the *A. tumefaciens* resuspension into falcon tubes containing freshly propagated calli (not older than two weeks) and incubated for 10 minutes at room temperature and mild agitation. After incubation, we placed the calli onto CIM supplemented with 100 µM acetosyringone and incubated them under callus induction conditions for two days. Finally, we washed the calli in SH-Medium supplemented with 300 mg L^-1^ cefotaxime and 250 mg L^-1^ ticarcillin for disinfection and placed them onto CIM plates supplemented with 300 mg L^-1^ cefotaxime and 250 mg L^-1^ ticarcillin (agroelimination-CIM). After three days of incubation, we collected the calli, washed them with distilled water, dried them on filter paper, snap them frozen in liquid nitrogen, and stored all the calli material at −80 °C for posterior quantification of metabolites.

### TYDC gene validation in S. polyrhiza calli

To validate the role of the *TYDC* and *TDC* candidate genes, we extracted free amino acids and amines. For this, we collected 10 mg of ground callus material and applied acidified methanol (MeOH:water:formic acid 15:4:1□v/v/v) without internal standards. After overnight incubation, we diluted an aliquot of the supernatant in an aqueous solution of isotope-labelled amino acids (algal amino acid mixture-13C-15N; Sigma-Aldrich) following the 1:100 proportion. After quantification of tyramine and tryptamine concentrations, we assessed the data with the linear model *Metabolite concentration∼construct*. The factor *construct* included wild type regenerated from calli, eGFP and empty vector, as well as *OsTYDC*, *SpTYDC1, SpTYDC2*, *SpTYDC3,* and *SpTDC*.

#### Transcriptome sequencing

To assess whether the identified tyrosine decarboxylases are transcriptionally regulated by the pre-treatment, we sequenced the transcriptomes of genotype Sp102 using the plant material from five replicates from each treatment and pre-treatment combination from the fitness and phenotype assay. To this end, we extracted total RNA from 10 mg of ground fresh plant material with the NucleoSpin RNA Plant Mini kit (Macherey-Nagel, Düren, Germany) following the manufacturer’s instructions. Then, library preparation with polyA enrichment and sequencing were performed by Novogene GmbH (Germany). After quality control, the libraries were sequenced on Illumina NovaSeq X Plus with 150 base pair paired end reads. From all 24 libraries, we obtained 3.8-9.9G; after filtering low-quality reads, these data resulted in an average of 38 Mio reads per library (Table **S4**). To quantify gene expression, Novogene performed a bioinformatic analysis of the data, which included quality control, mapping to the 7498-clone genome reference (*45*), gene expression quantification and differential expression between pre-treatments within control and aphid herbivory treatments. An average of 4×10^7^±8×10^6^ clean reads per library were obtained (Table **S4**) and mapped with HISAT2 (*52*) to the reference gene, obtaining an average mapping rate of 80%. For differential gene expression, abundance of transcripts–standardized in fragments per kilobase of transcript sequence per millions base pairs sequences (FPKM)–was used for hypothesis testing with DESeq2 (*53*), considering a negative binomial distribution and Wald test to obtain *P* values. To test whether *TYDCs* are differentially expressed, we considered *P*-values prior FDR corrections, as we assessed the expression values of only two genes.

### The role of tyramine in plant-aphid interaction

#### Chemical supplementation of tyramine to plants

To test whether tyramine alters plant and aphid fitness, as well as aphid melanization, we supplemented the plant growth medium with tyramine and measured its effects on aphid population growth rates and coloration. First, we determined which external tyramine concentrations produced *in planta* tyramine levels comparable to those observed in the indoor transgenerational experiment. Thereto, we dissolved tyramine (2-(4-Hydroxyphenyl)-ethylamine; Carl Roth, Karlsruhe, Deutschland) in DMSO and diluted it in N-medium to reach 0 µM, 100 µM, 200 µM and 400 µM tyramine and 0.05% v/v DMSO in N-medium. We then placed a single frond of genotype Sp102 into 150 mL of the tyramine-supplemented media, with five replicates per concentration. After eight days of free growth, we determined plant population growth rates as described above. Furthermore, we washed the plants by dipping them inside distilled water and subsequently determined shoot tyramine and tyrosine-derived amines following the amino acid extraction section.

Second, we examined whether supplementing plants with tyramine benefitted plant fitness under herbivory and reduced aphid population growth rates. Tyramine was dissolved in DMSO and added to N-medium to final concentrations of 0, 50, 100, 200, and 400 µM tyramine and 0.05% v/v DMSO in N-medium. To synchronize plant age, we grew plants of genotype Sp102 as single descendants for two generations inside polypropylene tubes. We then transferred a single frond—the first offspring of generation two—into 250 mL plastic beakers containing the tyramine-supplemented media. After five days of growth, we infested half of the beakers at each concentration with three adult aphids, approximately six days old, resulting in 12 infested and 12 un-infested replicates per concentration. After two and eight days, we counted the number of aphids to calculate daily per capita reproduction rates and population growth rates. Furthermore, we captured an image of the beakers at the beginning and end of the assay to estimate plant population growth rates. To assess whether tyramine supplementation altered plant fitness in the presence and absence of aphid herbivory, we fitted linear models with the mean plant population growth rates as a response variable and log-transformed tyramine concentration as well as treatment as explanatory variables. Log transforming tyramine concentration improved model fit. To assess whether tyramine supplementation reduced aphid fitness, we correlated the mean daily per capita reproduction rate with the external tyramine concentration using linear models.

#### Melanization in aphids

To assess whether tyramine affected aphid melanization, we first collected five wingless adult aphids per replicate, mounted them on sticky flea traps, and photographed with an MZ10 F Modular Stereo Microscope (Leica Microsystems, Wetzlar, Germany) using Leica Application Suite X v3.10.0.28982. Cuticle color intensity was quantified from the images with the “Histogram” function in ImageJ 64 v5 (*54*). To test whether tyramine reduces aphid redness—corresponding to increased melanization—we first averaged the pseudo-replicates within each replicate, then averaged across replicates, and finally correlated the resulting treatment means with tyramine concentration in a linear model. Second, we measure cuticle thickness through µCT. We collected five wingless adult individuals per replicate within 1mL of 4% paraformaldehyde in PBS (Thermo Fisher Scientific, Waltham, USA) and selected one random individual from the 0 µM and 400 µM tyramine treatments for the µCT. As a first step, we prepared the aphid samples. For this, we fixed the individuals in 4% paraformaldehyde in 80% ethanol for 48 h and after two subsequent 1h washing steps with 80% ethanol on a rocking shaker, we transferred the samples to 99% ethanol for 48 h for further dehydration. To contrast the soft tissue, we incubated the samples for 24 h in a freshly prepared 1% methanolic iodine solution, followed by three 1 h washing steps with 99% ethanol and three 1 h steps with 100% ethanol to remove all excess iodine. All steps were performed at room temperature. Once dehydrated, the samples were critically point dried in a CPD300 auto (Leica, Wetzlar, Germany) running on slowest possible settings for 35 cycles and medium fast heating at the end. Finally, we mounted the dried samples separately in an inverted 10 µl pipette tip and glued them to a sample holder of the µCT SkyScan 1272 (Bruker, Billerica, USA). We then scanned the aphid samples at 40 kV and 200 µA with an exposure time of 570 ms, rotational steps of 0.4°, and an isometric voxel resolution of 2.5 µm (image size 1640 × 930). Image stacks were reconstructed in NRecon (Bruker, Billerica, USA) and analyzed in ORS Dragonfly (Comet Technologies Canada Inc., Montreal, Canada). To estimate cuticle thickness, we first removed the pipet tip from the image stack and then created a region of interest (ROI) of the entire aphid body based on the histogram range. We filled the three dimensions within the ROI by iteratively applying dilate, close, and fill-inner-areas functions. The resulting solid ROI was inverted and used to overwrite all extra-aphid pixels to black. The resulting ROI was then inverted again, duplicated, and eroded to generate an ROI representing the aphid volume without the cuticle. We then subtracted the inner volume ROI from the total volume ROI to isolate the cuticle. Finally, to obtain the grey values of the cuticle, we inverted the cuticle ROI and applied it to a duplicated original image stack to obtain a new ROI via upper Otsu threshold setting. Finally, we extracted thickness meshes describing cuticle thickness across the whole aphid body.

#### Quantification of Buchnera

To quantify the load of *Buchnera* on *R. nymphaeae* fed on plants under different tyramine concentrations, we target the 16S rRNA and the *Ef1*α gene for normalization, reported for *R. maidis* and *R. padi* (*55*). We performed a quantitative PCR in four wingless adult individuals per tyramine treatment with the Rotor-Gene Q Real Time PCR(Qiagen, Venlo, Netherlands), using primers designed according to each symbiont 16S rRNA gene with amplification efficiency 93.8 and 99.6 % for *the* 16S rRNA and *Ef1*α gene, respectively (*56*). The qPCR reaction contained 10 µl of 2×KAPA SYBR® FAST (Roche, Basel, Switzerland), 0.4 µl of each primer, 1 ng of DNA, and Nuclease-free Water to reach 20 µl of total volume. The cycling conditions were 95 °C for 20 s, 40 cycles of 95 °C for 5 s and 60 °C for 20 s, followed by a melting curve. Considering four technical replicates per sample (N = 4), we first tested the efficiency of primers by dilution series of 0.1 ng, 1 ng and 10 ng purified DNA for each gene and measured DNA concentration with Qubit 4 Fluorometric Quantification system for ddDNA (Invitrogen). Then, we corrected for primer efficiency using the mathematical model of Pfaffl (*57*) for relative quantification in real-time PCR. We used linear models following the formula *fold change∼Treatment*.

#### Artificial diet supplemented with tyramine

To evaluate whether tyramine is sufficient to reduce aphid fitness and whether tyramine enhances cuticle redness, we measured per capita growth rates and color of aphids reared on sucrose solutions supplemented with ecologically relevant tyramine levels. A 35 % (w/v) sucrose solution in distilled water served as the base medium. Tyramine was first dissolved in 100% DMSO and then diluted to final concentrations of 0, 2, 5, 10, 20, 50, 100, 200, and 400 µM tyramine, each containing 0.5% v/v DMSO in the sucrose solution. We dispensed 8 mL of each solution into 60×15 mm Petri dishes (Sarstedt, Nümbrecht, Germany), with five replicates per concentration. To provide aphid a feeding site, we stretched Parafilm (Paul Marienfeld, Lauda Königshofen, Germany), cut it into 1 cm² squares, and floated three squares on the surface of the solution in each Petri dish. We then transferred eight similarly aged aphids, approximately 4 days old, onto the Parafilm in each dish and sealed the plates with Parafilm. Five days later, we counted the aphids to calculate per capita growth rates. We then collected five wingless adults per replicate for color analysis as described above. After averaging per capita growth rates and aphid redness for each treatment, we used linear models to correlate these means with the natural logarithm of tyramine concentration. The log transformation improved model fit.

#### Stable transformation and regeneration of *S. polyrhiza* TYDC lines

To obtain stable transgenic *S. polyrhiza* lines containing the SpTYDC constructs, as well as the controls eGFP and empty vector, we incubated the transformed calli for two weeks on agroelimination-CIM plates and then transferred the calli to low selection medium composed of agroelimination-CIM containing 5 mg L^-1^ hygromycin. After two weeks of incubation under callus induction growth conditions, we transfer the resistant calli to strong selection medium, composed of agroelimination-CIM containing 10 mg L^-1^ hygromycin and incubated the calli for two weeks, allowing only the propagation of transformed calli and eliminating any residual wild-type calli.

To regenerate the plants, we follow the steps of Barragán-Borrero(*25*). Briefly, we fragmented the transformed calli and placed small pieces (ca. 3 mm) on frond regeneration medium (FRM) composed of MS-Medium including vitamins (Duchefa) supplemented with, 1% (w/v) sucrose, 500 mg L^-1^ 2-morpholinoethanesulfonic acid monohydrate, 2.2 mg L^-1^ citric acid monohydrate, 100 mg L^-1^ polyvinylpyrrolidone average mol wt 40000, 1 mg L^-1^ thidiazuron, and 0.4% (w/v) gelrite, pH 5.8, and incubate within long day conditions (16:8h light/dark), 20 μmol photons m^−2^ s^−1^ and 26 °C. After four weeks of incubation, leaf-like structures were formed, which we selected and transferred to liquid SH-Medium containing 1% (w/v) sucrose and cultured within long day conditions (16:8h light/dark), 85 μmol photons m^−2^ s^−1^ and 26 °C until full-regenerated colonies were formed (ca. two weeks). Finally, we acclimated the regenerated plants to SH-Medium without sucrose and further propagated the plants within long day conditions (16:8 h light/dark), 135 μmol photons m^−2^ s^−1^ and 26 °C.

#### Genotyping of transgenic lines

To genotype the transformed plants, we first screened the regenerated lines using Thermal Asymmetric Interlaced PCR (TAIL-PCR)(*58*) and through metabolite profiling (Fig. **S4F**). We then selected independent lines from each construct (SpTYDC1, SpTYDC2, eGFP and empty vector) to confirm their independence and determine transgene copy number using restriction fragment length polymorphism (RFLP) analysis via Southern blotting (Fig. **S4G**). To obtain DNA for TAIL-PCR and RFLP, we snap-froze fronds from regenerated plants in liquid nitrogen and ground the frozen material to a fine powder using a grinding mill (RETSCH MM 400 Mixer Mill). Then, we extract the total DNA using the cetyltrimethylammonium bromide (CTAB) extraction method (*59*), including treatment with RNAse A (Sigma-Aldrich), and quantified using a Thermo Scientific NanoDrop spectrophotometer.

For the initial screening of lines, we carried out a TAIL-PCR in the TYDC lines and included as controls, the regenerated wild type and the empty vector line #1. We used an arbitrary degenerate primer (5’-ntcgastwtsgwgtt-3’) and three sequence-specific primers which anneal to the NOS terminator sequence (5’-atgattagagtcccgcaattatac-3’; 5’-gcaaactaggataaattatcgcgc-3’; and 5’-ggtcgacgcgtcaattgatatc-3’). We used the DreamTaq DNA Polymerase (Thermo Scientific™) protocol to conduct the PCR reactions. The amplification products were then separated and detected via agarose gel electrophoresis. Next, we measured the concentrations of the amines tyramine, L-Dopa, dopamine and octopamine, as well as the amino acid L-tyrosine. We cultivated the SpTYDC1, SpTYDC2, wild type, eGFP and empty vector lines for seven days within 250 mL transparent polypropylene beakers filled with 150 mL 0.5 SH-medium and covered with perforated transparent lids. Then, we harvest the shoots of the 12 replicates per line and flash-froze the material for posterior metabolite extraction using the amino acid method. To assess the data, we used the model *Metabolite ∼Line +(1|Construct)+(1|Batch)*. With the results, we select potential TYDC independent lines for RFLP analysis.

For RFLP analyses, we treated 20 μg total DNA with the restriction enzyme HindII. The resulting fragments were then resolved by electrophoresis using 1% (w/v) agarose gel. Before blotting, DNA denaturation was conducted by treating the gel sequentially with (i) 0.25 M HCl for 15 min, then rinsed with distilled water; (ii) 0.5 M NaOH for 30 min, then rinsed with distilled water; (iii) 0.5 M NaOH and 1.5 M NaCl for 30 min, then rinsed with distilled water; (iv) and lastly incubated in 1 M Tris, 3 M NaCl (pH 6.5) for 15 min. Next, we transferred the DNA from the gel to a positively charged nylon membrane (Cytiva Life Sciences™ Amersham™ Hybond-N+ Membrane) via capillary blotting using 10 × SSC buffer (1.5 M NaCl, 0.15 M trisodium citrate), and UV-crosslinked twice (Boekel scientific, Philadelphia, USA) at 0.12 J cm^-2^. For hybridization, we prepared DNA probes using the DIG-High Prime DNA Labeling and Detection kit (Roche), following the manufacturer’s instructions. We chose as a template to produce a labelled probe suitable for the genotyping of all constructs, the selectable marker sequence (hptII gene), which was amplified via PCR (Phusion™ High-Fidelity DNA Polymerase; Invitrogen) using the primer pair 5’-caattagagtctcatattcactctcaa-3’ and 5’-ctttattgccaaatgtttgaacgatc-3’. After overnight hybridization at 42 °C in DIG Easy Hyb buffer (Roche) containing 10 ng ml^-1^ of DIG-labelled probe, we washed the membrane sequentially with (i) 2 × SSC buffer with 0.1% (w/v) sodium dodecyl sulphate (SDS) for 20 min at room temperature; (ii) 0.5 × SSC buffer with 0.1% (w/v) SDS for 15 min at 68 °C twice; (iii) 0.1 M maleic acid, 0.15 M NaCl, 0.3% (v/v) tween 20, and pH 7.5 for 5 min at room temperature. Then, we incubated the membrane for 1 h at room temperature with Blocking Solution (Roche), 1 h at room temperature with 150 mU ml^-1^ Anti-Digoxigenin-AP (Roche) diluted in Blocking Solution, and performed a final washing by incubating the membrane twice for 15 min at room temperature with 0.1 M maleic acid, 0.15 M NaCl, 0.3% (v/v) tween 20. Before detection, we first equilibrated the membrane for 5 min in detection buffer (0.1 M Tris and 0.1 M NaCl, pH 9.5) and then performed the color detection by overnight incubation with 20µl ml^-1^ NTB/BCIP stock solution (Roche) diluted in detection buffer. To document the color development, we used the GelDoc Go Gel Imaging System (Bio-Rad, Hercules, USA).

#### Bioassays with TYDC transgenic lines

To assess whether endogenous production of tyramine alters the resistance of *S. polyrhiza* to *R. nymphaeae*, we measured plant and aphid reproduction when aphids were added to *SpTYDC1 and SpTYDC2* lines, lines that constitutively overexpress the tyrosine decarboxylases. We pre-cultivated regenerated wild type Sp162, eGFP#1, eGFP#2, eGFP#3, empty vector#1, SpTYDC1#2, SpTYDC1#6, SpTYDC2#4, SpTYDC2#5 and SpTYDC2#9 plants within autoclaved 2.6 L boxes (Lock & Lock) filled with 300 mL of 0.5 SH-medium for two weeks under 26 °C, 135 µmol photons m^-2^ s^-1^, 16:8 h light/dark cycle. At the same time, we synchronized the age of aphids by placing adult aphids on wild type plants within autoclaved 2.6 L boxes (Lock & Lock) filled with 300 mL of 0.5 SH-medium. After 24 h we removed the adults and let the nymphs mature for six days before moving them to the transgenic plants.

When setting up the assay, we adjusted the number of fronds at the start of the assay to achieve a similar plant surface area for each line, thereby resulting in comparable herbivore pressure and food availability for the aphid. The adjustments were needed since transgenic plants were smaller than the wild type. Fronds of each line were placed separately within 250 mL transparent polypropylene beakers filled with 150 mL 0.5 SH-medium and covered with perforated transparent lids, with 12 replicates per line. To obtain similar surface area for each line, we first cut 1 cm^2^ out from the bottom of a polypropylene beaker and used the hole as reference to cover each beaker with a similar plant surface area. This means that we used different number of starting fronds. Then, we estimated the surface area with ImageJ 64 v5 (*54*) and added or removed fronds if necessary. In average, we started with five wild type fronds and 12 transgenic fronds per beaker. The experiment was initiated over three consecutive days, using a subset of each line per day to make handling feasible.

To start the assays, we introduced three six-day synchronized adult aphids per beaker and let both plants and aphids reproduce freely for seven days. To measure aphid performance, we counted the number of adult aphids and nymphs, distinguishing between dead and alive, after seven days of free reproduction. To quantify surface area and frond number, we captured images at the beginning and end of each assay using a camera box installation with a webcam (HD Pro Webcam C270; webcam software 2.12.8). We analyzed surface area of plants with ImageJ 64 v5 (*54*) and counted fronds with dotdotGoose v1.5.3 (*47*). Plant material was stored at −20 °C and subsequently used to quantify tyramine accumulation, as described above in the free amino acids and amines extraction section.

To analyze whether TYDC overexpression alters plant fitness, we estimated plant population growth rates as described above. We used the mixed-effects models *Plant fitness ∼Line +(1|Construct)+(1|Batch)* and *Plant fitness ∼Gene+(1|Construct/Line)+(1|Batch)*.

To assess whether *in planta* tyramine overaccumulation alters aphid performance, we first estimated nymph and total aphid (all aphid stages) death rate on the different constructs. As death rates follow a negative binomial distribution, we used the mixed-effects model *Number of dead aphids ∼Line +(1|Construct)+(1|Batch), offset =log(exposure)*, where “Batch” represents the group of plants tested per day and exposure refers to density-time at risk as show in formula (**3**). Similarly, to assess the death rate differences between the control group and the SpTYDC plants we used the model *Number of dead aphids ∼Gene+(1|Construct/Line)+(1|Batch), offset =log(exposure)*. To test whether TYDC overexpression alters the reproductive rate of aphids, we used the model *Aphid Fitness∼Line +(1|Construct)+(1|Batch)* and *Aphid fitness ∼Gene +(1|Construct/Line)+(1|Batch)*. To verify the repeatability of our results, we performed a second experiment using a subset of the genotypes, including wild type (regenerated Sp162), eGFP#3, empty vector, SpTYDC1#2, SpTYDC1#6, SpTYDC2#4, SpTYDC2#5 and SpTYDC2#9 plants, using the same conditions and data analysis as described above.

### The role of jasmonates in plant-aphid interaction

#### External application of methyl-jasmonate to plants

To evaluate the impact of jasmonates on aphid reproduction, we monitored aphid population growth rates on Sp102 plants exposed to different levels of methyl-jasmonate. As methyl-jasmonate (Duchefa Biochemie) is volatile, we executed two different approaches: short-term and continuous methyl-jasmonate induction. For both approaches, we first synchronized plant age by propagating plants as single descendants for two generations.

For the short-term induction, we transferred the first offspring of generation three into 30 mL polypropylene tubes (Fisher Scientific) containing N-medium with different levels of methyl-jasmonates and 0.5 % v/v DMSO. Control N-medium contained only 0.5% v/v DMSO. After three days of induction on the single fronds, we moved the colony—the original frond together with its offspring—into 250 mL transparent polypropylene beakers filled with 150 mL N-medium and introduced three six-day synchronized aphids to half of the beakers. We counted the number of nymphs after 24 h or 48 h and stored the plant shoots at −20 °C for posterior metabolite extraction following the targeted screening method.

We performed the short-term induction experiment twice: first using eight replicates with 0 µM, 20 µM, 40 µM, 60 µM, 80 µM and 100 µM methyl-jasmonates and then 16 replicates with 0 µM, 25 µM, 50 µM, 100 µM, 150 µM and 200 µM methyl-jasmonate. To assess whether methyl-jasmonate affects aphid performance in a dose-dependent manner, we fitted the model *Mean performance ∼ natural spline of MeJA concentration*, considering two and three degrees of freedom for the first and the second experiment, respectively. MeJA refers to methyl-jasmonate. We used a natural spline to model the non-linear relationship while keeping the curve linear at the extremes. To assess the effects of 60 µM and 80 µM methyl jasmonate on aphid performance, we fitted the per capita growth rate into the mixed-effects model *Performance ∼MeJA +(1|Replicate)*. For the continuous induction, we placed the first *S. polyrhiza* offspring of generation three inside 150 mL of N-medium containing different levels of methyl-jasmonate within transparent polypropylene beakers covered with perforated transparent lids. After 72 h, we introduced three six-day synchronized aphids to half of the beakers, each containing now a plant colony—the original frond together with its offspring—and let both plants and aphids reproduce for eight days. To estimate aphid and plant fitness, we counted the number of adult aphids and total aphids at the end of assays and took pictures of the plants at the beginning and end of assays, as described above. We conducted this experiment twice: first using 0 µM, 25 µM, 50 µM, 100 µM, 150 µM and 200 µM methyl-jasmonate with 16 replicates per treatment and second using 0 µM and 80 µM methyl-jasmonate with 12 replicates per treatment. In the second assay, we excluded four out of 12 replicates because winged morphs appeared by day two, strongly reducing reproduction. To test whether methyl-jasmonate altered aphid performance though per capita growth rate (adult aphids relative to the total number of aphids), we fitted the model *Mean fitness∼natural spline of MeJA concentration*, with three degrees of freedom. Similarly, to assess the protective role of jasmonates on plants, we used the model *Mean fitness ∼natural spline of MeJA concentration *Herbivory treatment*, with three degrees of freedom. Finally, to test the differences in aphid fitness (daily population growth rates) between 0 µM and 80 µM of continuous methyl-jasmonate we fitted the model *Aphid fitness ∼MeJA concentration +(1|Batch)*.

To identify the effects that increased jasmonates have on the levels of free amino acids, we collected plant tissue after three days of induction. As free animo acids, we quantified the concentrations of L-Alanine, L-Arginine, L-Asparagine, L-Aspartic acid, L-Glutamic acid, L-Glutamine, L-Isoleucine, L-Leucine, L-Lysine, L-Phenylalanine, L-Proline, L-Serine, L-Threonine, L-Tryptophane, L-Valine, Ornithine, Glycine, L-Histidine, L-Methionine, L-Tyrosine, Citrulline and GABA. Then, we calculated the fold change per amino acid between treated and control plants and fitted the model *natural logarithm of fold change ∼MeJA concentration +(1|Amino acid) +(1|Lineage)*.

### Transgenerational experiments in natural conditions

To assess whether plants inherit stress responses outdoors, we used an outdoor mesocosm experiment at the Swiss Federal Institute of Aquatic Science and Technology (Eawag) in Dübendorf, Switzerland, in which genotype Sp102 had been grown for two consecutive years with and without herbivory of the *R. nymphaeae* genotype Rn001 (pre-treatment or pond treatment) in 2021 and 2022 (*27*). Ponds had a size of 4×4 m, depth of 1.5 m and volume of around 15 m^3^, and at the peak of duckweed population growth, contained approximately 1 million duckweed individuals.

#### Phenotypic plasticity within a year

To test whether tyramine and jasmonates are phenotypically plastic outdoors, we randomly selected plants of the ponds in July and August 2021. Plant material of whole fronds was cleared of aphids, rinsed in tap water, briefly dried with a tissue paper, flash-frozen in liquid nitrogen and subsequently stored at −80 °C until metabolite analysis, see below. To measure the levels of these metabolites in turions, we collected detached turions from the bottom of the ponds in the shallow water zones in October 2021. Turions were briefly dried with tissue paper, flash-frozen in liquid nitrogen and stored at −80 °C until metabolite analysis.

For metabolite quantification, we freeze-dried shoot and turion material, ground it to a fine powder, and extracted approximately 10 mg freeze-dried material for tyramine and jasmonate analysis following the protocol used in the indoor transgenerational experiment. To test whether the pond treatment alters metabolite accumulation, we fitted mixed-effect models with pond treatment as a fixed effect and group of ponds as a random effect: *Metabolite ∼Pond treatment +(1|Group of ponds)*, where “Group of ponds” represented the physical four-pond block in the field (Fig **6A**).

#### Inheritance of stress responses across years

To assess whether tyramine and jasmonates are transgenerationally plastic across years, and thereby alter plant and aphid fitness, as well as plant morphology, we performed *in situ* transplant experiments. During spring of 2022 (calendar week 17, Fig **6B**), we collected 20 *S. polyrhiza* surface-floating turions per pond and transferred them to boxes (17 cm width ×12 cm deep ×11.5 cm high, Eurobox, Auer Packaging) that were floating within the aphid herbivory ponds. The lids of the boxes were modified with a mesh to allow natural light and air exchange, but restricted aphid from scaping the box. Thus, each herbivory pond contained two floating boxes: one held its own turions, and the other held turions collected from a neighboring control pond. We used only the herbivory ponds to avoid contaminating the control ponds. After three weeks of free growth (“recovery”), we transferred 20 newly emerged fronds from each box into fresh floating boxes and grew the plants freely for another three weeks under control conditions or *R. nymphaeae* herbivory, with 5 aphids per box (“treatment”). We then harvested the plants: we counted the aphids, gently dried the plants with tissue paper and flash-froze the plant material in liquid nitrogen. All plant material was stored at −80 °C until metabolite extraction. We quantified plant surface area and frond number from photographs taken at the start and end of the treatment phase with a Nikon D5300 (Nikon, Tokyo, Japan) mounted in a camera box. Surface area was measured in ImageJ 64 v5 (*54*) and fronds counted with dotdotGoose v1.5.3 (*47*).

To quantify tyramine and jasmonate concentrations, we ground the frozen plant and turions material, aliquoted 20 mg ground material and subsequently measure tyramine and jasmonate levels as described above. To assess whether the pre-treatment altered metabolite accumulation in turions, we fitted mixed-effects models with the ancestral treatment as fixed effect and the group of four ponds as random factor. Therefore, the model was *Metabolite ∼Pre-treatment +(1|Receiving pond)*. To assess whether the pre-treatment altered metabolite accumulation in plants, we fitted mixed-effect models with the pre-treatment and treatment as fixed effects and the receiving pond as random effect. The model was: *Metabolite ∼Pre-treatment *Treatment +(1|Receiving pond)*. Plant fitness and morphology were analyzed in a similar way, using the model *Variable ∼Pre-treatment *Treatment +(1|Receiving pond)*, whereas we used the model *Variable ∼Pre-treatment +(1|Receiving pond)* for aphid fitness.

#### Inheritance of stress responses within a year

To test whether plants are transgenerationally plastic within a year, we performed a similar reciprocal transplant experiment. In July (calendar week 30), when aphid intensities almost peaked (Fig **6B**), we collected 40 adult fronds carrying a offspring frond from each pond. We brushed off all aphids, marked both the parent and its offspring with a permanent marker (Stabilo OHPen Universal, Heroldsberg, Germany), and placed 20 marked parents carrying one offspring into the floating boxes positioned either in their home pond or in a neighboring pond of the opposite treatment. Thus, each pond contained floating boxes with plants from both treatments.

After three weeks of free growth without aphids (“recovery”), we took 40 unmarked fronds from each box, split them equally between two new boxes floating in the same pond, and introduced five adult aphids to one of the boxes; the other served as a control. After a further three-week free growth period (“treatment”), we counted aphids and determined surface area growth rates and plant morphology as described above. We then flash-froze shoots in liquid nitrogen and stored at −80 °C until metabolite extraction. To quantify metabolites, we ground the frozen plant material and used 20 mg ground material to determine tyramine and jasmonate levels as described above. To assess whether the pre-treatment altered metabolite accumulation, we fitted mixed-effect models with the pre-treatment and treatment as fixed effects and the initial pond and receiving pond as random effects. “Initial pond” is the pond from which the plants were taken before the recovery phase; “Receiving pond” is the ponds in which the recovery and treatment were performed: *Metabolite ∼Pre-treatment *Treatment +(1|Initial pond)+(1|Receiving pond)*. For plant fitness and morphology traits, we accounted for the herbivory or control conditions the ponds had at the start of the fitness and phenotype assays, yielding: *Variable ∼Pre-treatment *Treatment +(1|Initial pond)+(1|Receiving pond)+(1|Pond treatment)*. For aphid fitness, we included as random factor the number of adults fronds (11 to 15 adult fronds) when starting fitness and phenotype assays, as the developmental stage of the plant may affect aphid performance, using the model *Variable∼Pre-treatment +(1|Initial pond)+(1|Receiving pond)+(1|Pond treatment)+(1|Adult fronds)*. For all analysis, we excluded replicates in which the control treatment got contaminated with aphids or were aphids died due to fungi infection.

### Transgenerational experiments after turion formation

To assess whether turion formation erases the transgenerational increase in tyramine concentrations, we induced turion formation in *S. polyrhiza* genotype Sp102 indoors. After the eight lineages from the indoor transgenerational experiment reached generation seven—five generations of herbivory pre-treatment and two generations under recovery phase—we moved half of the first offspring and half of the second offspring within 30 mL polypropylene tubes (Fisher Scientific) filled with 25 mL of low-phosphate N-medium—2 µM KH_2_PO_4_ instead of 150 µM KH_2_PO_4_ used in normal N-medium (*24*). Then, we cultivated the plants under 18 °C, 150 µmol photons m^-2^ s^-1^ and 16:8 h light/dark cycle until each lineage of generation eight, but not nine or 10, produced the first turion.

To continue with the single descendant propagation, we collected the turions—current generation nine—if they easily detach from the parental plant when touching the surface of the turion. Then, we softly dried the turions on tissue paper and gently placed each of them in one well of a six-well plate (Fisher scientific) filled with 10 mL of N-medium. We covered the plates with their corresponding lids (Fisher scientific) and placed the plates at 28 °C, 150 µmol photons m^-2^ s^-1^ and 16:8 h light/dark cycle. To increase germination efficiency, we gently placed the dry turions on the surface of the medium, with the same side of the turion facing the medium as when the turion was generated in the parental plant. Once the new plants germinated—generation 10, we moved the plants into 30 mL polypropylene tubes (Fisher Scientific) filled with 25 mL of N-medium. Finally, we placed the offspring (generation 11) into control conditions or recurring aphid herbivory for the fitness and phenotype assays and collected the shoots of plants for posterior metabolite quantification.

To measure the concentrations of tyramine in plant tissue, we followed the screening method described above. To assess the differences in tyramine concentration among treatments, we fitted the model *Tyramine ∼Pre-treatment *Treatment +(1|Lineage)*.

## Acknowledgments

We would like to acknowledge Anne Schreyer, Marieke Theiner, Sofie Rutenbeck, Barbara Eppard and Büsra Karagöz for executing the external application of methyl-jasmonate and tyramine experiments, the artificial diet experiments and quantifying the metabolites from the indoor experiments. As well, we thank Erica Venberg and Jessica Nietulski for performing the experiments with transgenic plants. We thank Wolf Frommer for providing the rice seeds and Arturo Mari-Ordoñez for sharing with us the binary vector CP101 (eGFP). As well, we want to acknowledge the support of Marie Serwaty-Sárazová and Sara Noure in the field bioassays, Christoph Walcher, Christoph Vorburger and Piet Spaak for facilitating us the work at EAWAG, and Holger Schön and David Martín Fernandez from the University of Münster for constructing the floating boxes for the outdoor experiments and the lids for our indoor bioassays.

## Data, code, and materials availability

All data and code needed to evaluate and reproduce the results in the paper are present on Zenodo https://zenodo.org/records/20144931 and on NCBI under the project PRJNA1405110 https://dataview.ncbi.nlm.nih.gov/object/PRJNA1405110.

## REFERENCES

1. R. Bonduriansky, T. Day, Nongenetic inheritance and its evolutionary implications. Annu. Rev. Ecol. Evol. Syst. 40, 103 (2009).

2. R. E. O’Dea, D. W. A. Noble, S. L. Johnson, D. Hesselson, S. Nakagawa, The role of non-genetic inheritance in evolutionary rescue: epigenetic buffering, heritable bet hedging and epigenetic traps. *Environ*. Epigenetics 2, dvv014 (2016).

3. A. Chávez et al., Copper-induced transgenerational plasticity in plant defence boosts aphid fitness. Plant Cell Environ. 48, 3997 (2025).

4. A. Grafen, in Reproductive Success: Studies of Individual Variation in Contrasting Breeding Systems, T. H. Clutton-Brock, Ed. (University of Chicago Press, 1998).

5. S. H. Chung et al., Herbivore exploits orally secreted bacteria to suppress plant defenses. PNAS 110, 15728 (2013).

6. C. Brütting et al., Cytokinin transfer by a free-living mirid to Nicotiana attenuata recapitulates a strategy of endophytic insects. eLife 7, e36268 (2018).

7. G. Gibson, The environmental contribution to gene expression profiles. Nat. Rev. Genet. 9, 575 (2008).

8. D. Anastasiadi, C. J. Venney, L. Bernatchez, M. Wellenreuther, Epigenetic inheritance and reproductive mode in plants and animals. Trends Ecol. Evol. 36, 1124 (2021).

9. P. Crevillén et al., Epigenetic reprogramming that prevents transgenerational inheritance of the vernalized state. Nature 515, 587 (2014).

10. B. Pasaribu et al., Genomics of turions from the greater duckweed reveal its pathways for dormancy and re-emergence strategy. New Phytol. 239, 116 (2023).

11. U. Grossniklaus, B. Kelly, A. C. Ferguson-Smith, M. Pembrey, S. Lindquist, Transgenerational epigenetic inheritance: how important is it? Nat. Rev. Genet. 14, 228 (2013).

12. M. Huber, A. Chávez, Assessing rapid adaptation through epigenetic inheritance: a new experimental approach. Plant Cell Environ. 48, 1494 (2025).

13. A. Chávez, M. Schäfer, I. Finkemeier, S. Xu, M. Huber, Heritable transgenerational fitness variation correlates with copper resistance in the clonal duckweed Spirodela polyrhiza. Proc. R. Soc. B, (2026).

14. M. Erb, Plant defenses against herbivory: closing the fitness gap. Trends Plant Sci. 23, 187 (2018).

15. S. Xu et al., Low genetic variation is associated with low mutation rate in the giant duckweed. Nat. Commun. 10, 1243 (2019).

16. P. Ziegler, K. Adelmann, S. Zimmer, C. Schmidt, K.-J. Appenroth, Relative in vitro growth rates of duckweeds (Lemnaceae) – the most rapidly growing higher plants. Plant Biol. 17, 33 (2015).

17. E. Landolt, Biosystematic investigation in the family of duckweeds (“Lemnaceae”) (vol.2). The family of Lemnaceae - a monographic study. Veröffentlichungen des Geobotanischen Institutes der Eidg. Tech. Hochschule, Stiftung Rübel, in Zürich 1, 567 (1986).

18. T. Hance, D. Nibelle, P. Lebrun, G. van Impe, C. van Hove, Selection of Azolla forms resistant to the water lily aphid *Rhopalosiphum nymphaeae. Susceptibility of* Azolla *forms to* Rhopalosiphum nymphaeae. Entomol. Exp. Appl. 70, 19 (1994).

19. A. K. Hansen, N. A. Moran, Aphid genome expression reveals host–symbiont cooperation in the production of amino acids. PNAS 108, 2849 (2011).

20. Y. Chen et al., An aphid RNA transcript migrates systemically within plants and is a virulence factor. PNAS 117, 12763 (2020).

21. A. Korgaonkar et al., A novel family of secreted insect proteins linked to plant gall development. Curr. Biol. 31, 1836 (2021).

22. C. Sempruch et al., Influence of selected plant amines on probing behaviour of bird cherry-oat aphid (*Rhopalosiphum padi* L.). Bull. Entomol. Res. 106, 368 (2016).

23. T. Herben, J. Klimešová, Evolution of clonal growth forms in angiosperms. New Phytol. 225, 999 (2020).

24. K. J. Appenroth, S. Teller, M. Horn, Photophysiology of turion formation and germination in *Spirodela polyrhiza*. Biol. Plant 38, 95 (1996).

25. V. Barragán-Borrero et al., Strain, procedures, and tools for reproducible genetic transformation and genome editing of the emerging plant model *Spirodela polyrhiza*. New Phytol. 250, 735 (2026).

26. Y. Wang, S. Xu, A high-quality genome assembly of the waterlily aphid *Rhopalosiphum nymphaeae*. Sci. Data 11, 194 (2024).

27. M. Schäfer, et al., Aphid herbivory on macrophytes drives adaptive evolution in an aquatic community via indirect effects. PNAS 122, e2502742122 (2025).

28. R Core Team, R: a language and environment for statistical computing. (2024).

29. R. Hunt, Plant growth curves. The functional approach to plant growth analysis. (Edward Arnold Ltd, London, 1982).

30. J. R. Carey, Applied demography for biologists: special emphasis on insects. (Oxford University Press, 1993).

31. J. H. Matis, T. R. Kiffe, T. I. Matis, D. E. Stevenson, Stochastic modeling of aphid population growth with nonlinear, power-law dynamics. Math. Biosci. 208, 469 (2007).

32. M. Huber, S. Gablenz, M. Höfer, Transgenerational non-genetic inheritance has fitness costs and benefits under recurring stress in the clonal duckweed *Spirodela polyrhiza*. Proc. R. Soc. B 288, 20211269 (2021).

33. M. E. Brooks et al., glmmTMB balances speed and flexibility among packages for zero-inflated generalized linear mixed modeling. The R Journal 9, 378 (2017).

34. P. J. Aphalo, ggpmisc: Miscellaneous extensions to ‘ggplot2’. (2024).

35. R. V. Lenth, emmeans: estimated marginal means, aka least-squares means. (2024).

36. H. Torsten, F. Bretz, P. Westfall, Simultaneous inference in general parametric models. Biom. J. 50, 346 (2008).

37. F. Hartig, DHARMa: residual diagnostics for hierarchical (multi-level / mixed) regression models. (2022).

38. J. Fox, S. Weisberg, Visualizing fit and lack of fit in complex regression models with predictor effect plots and partial residuals. J. Stat. Softw. 87, 1 (2018).

39. H. Wickham, ggplot2: Elegant graphics for data analysis. Use R! (Springer New York, ed. 2, 2016), pp. 2197–5736.

40. H. Wickham, J. Bryan, readxl: read excel files. (2023).

41. T. Barrett et al., data.table: extension of ‘data.fram’. (2024).

42. J. Silge, D. Robinson, tidytext: text mining and analysis using tidy data principles in R. J. Open Source Softw. 1, 37 (2016).

43. H. Wickham, R. François, L. Henry, K. Müller, D. Vaughan, dplyr: a grammar of data manipulation. (2023).

44. A. Malacrinò et al., Induced responses contribute to rapid adaptation of *Spirodela polyrhiza* to herbivory by *Lymnaea stagnalis*. *Commun*. Biol. 7, 81 (2024).

45. Y. Wang et al., Population genomics and epigenomics of *Spirodela polyrhiza* provide insights into the evolution of facultative asexuality. *Commun*. Biol. 7, 581 (2024).

46. T. Olivoto, Lights, camera, pliman! An R package for plant image analysis. Methods Ecol. Evol. 13, 789 (2022).

47. P. J. Ersts, DotDotGoose.

48. D. An et al., Plant evolution and environmental adaptation unveiled by long-read whole-genome sequencing of Spirodela. PNAS 116, 18893 (2019).

49. T. A. Hall, BioEdit: a user-friendly biological sequence alignment editor and analysis program for Windows 95/98/NT.. Nucleic Acids Symp. Ser. 41, 95-98 (1999).

50. S. Park et al., Induced tyramine overproduction in transgenic rice plants expressing a rice tyrosine decarboxylase under the control of methanol-inducible rice tryptophan decarboxylase promoter. Bioprocess Biosyst. Eng. 35, 205 (2012).

51. W. D, G. J., Transformation of Agrobacterium using the freezethaw method. Cold Spring Harb. Protoc. 7, (2006).

52. A. Mortazavi, B. A. Williams, K. McCue, L. Schaeffer, B. Wold, Mapping and quantifying mammalian transcriptomes by RNA-Seq. Nat. Methods 5, (2008).

53. M. I. Love, W. Huber, S. Anders, Moderated estimation of fold change and dispersion for RNA-seq data with DESeq2. Genome Biol. 15, 550 (2014).

54. C. A. Schneider, W. S. Rasband, K. W. Eliceiri, NIH Image to ImageJ: 25 years of image analysis. Nat. Methods 9, 671 (2012).

55. S. Liu et al., Effects of host plants on aphid feeding behavior, fitness, and *Buchnera aphidicola* titer. Insect Sci. 32, 927 (2025).

56. S. Liu et al., Secondary symbionts affect aphid fitness and the titer of primary symbiont. Front. Plant Sci. Volume 14-2023, (2023).

57. M. W. Pfaffl, A new mathematical model for relative quantification in real-time RT–PCR. Nucleic Acids Res. 29, e45 (2001).

58. Y.-G. Liu, Y. Chen, High-efficiency thermal asymmetric interlaced PCR for amplification of unknown flanking sequences. BioTechniques 43, 649 (2007).

59. J. J. Doyle, D. J. L., Isolation of Plant DNA From Fresh Tissue. (Focus, Amsterdam, Netherlands, 1990), vol. 12.

60. J.-J. Xu, X. Fang, C.-Y. Li, L. Yang, X.-Y. Chen, General and specialized tyrosine metabolism pathways in plants. aBIOTECH 1, 97 (2020).

61. K. Brandau, J. Axelrod, The biosynthesis of octopamine. Naunyn-Schmiedeb. Arch. Pharmacol 273, 123 (1972).

62. A. R. Soares, et al., The role of L-DOPA in plants. Plant Signal. Behav. 9, e28275 (2014).

63. T. Lehmann, S. Pollmann, Gene expression and characterization of a stress-induced tyrosine decarboxylase from *Arabidopsis thaliana*. FEBS Letters 583, 1895 (2009).

64. K. Lee, K. Kang, M. Park, S. Park, K. Back, Enhanced octopamine synthesis through the ectopic expression of tyrosine decarboxylase in rice plants. Plant Sci. 176, 46 (2009).

65. P. J. Facchini, V. De Luca, Differential and tissue-specific expression of a gene family for tyrosine/dopa decarboxylase in opium poppy. J. Biol. Chem. 269, 26684 (1994).

66. P. J. Facchini, V. De Luca, Phloem-specific expression of tyrosine/dopa decarboxylase genes and the biosynthesis of isoquinoline alkaloids in opium poppy. Plant Cell 7, 1811 (1995).

67. P. Kawalleck, H. Keller, K. Hahlbrock, D. Scheel, I. E. Somssich, A pathogen-responsive gene of parsley encodes tyrosine decarboxylase. J. Biol. Chem. 268, 2189 (1993).

68. J. Günther et al., Separate pathways contribute to the herbivore-induced formation of 2-phenylethanol in poplar. Plant Physiol., pp.00059.2019 (2019).

69. M. López-Meyer, C. L. Nessler, Tryptophan decarboxylase is encoded by two autonomously regulated genes in *Camptotheca acuminata* which are differentially expressed during development and stress. Plant J. 11, 1167 (1997).

70. C. Qiao et al., Functional characterization of a catalytically promiscuous tryptophan decarboxylase from camptothecin-producing *Camptotheca acuminata*. Front. Plant Sci. 13 (2022).

71. V. De Luca, C. Marineau, N. Brisson, Molecular cloning and analysis of cDNA encoding a plant tryptophan decarboxylase: comparison with animal dopa decarboxylases. PNAS 86, 2582 (1989).

72. Y. Yamazaki, H. Sudo, M. Yamazaki, N. Aimi, K. Saito, Camptothecin biosynthetic genes in hairy roots of *Ophiorrhiza pumila*: cloning, characterization and differential expression in tissues and by stress compounds. Plant Cell Physiol. 44, 395 (2003).

73. S. G. Kang, H. K. Jeong, E. Lee, S. Natarajan, Characterization of a lipoate-protein ligase A gene of rice (*Oryza sativa* L.). Gene 393, 53 (2007).

74. W. Liu et al., Tryptophan decarboxylase plays an important role in ajmalicine biosynthesis in *Rauvolfia verticillata*. Planta 236, 239 (2012).

75. M. Gutensohn et al., Role of aromatic aldehyde synthase in wounding/herbivory response and flower scent production in different Arabidopsis ecotypes. Plant J. 66, 591 (2011).

76. Y. Kaminaga et al., Plant phenylacetaldehyde synthase is a bifunctional homotetrameric enzyme that catalyzes phenylalanine decarboxylation and oxidation *J*. Biol. Chem. 281, 23357 (2006).

77. T. Koeduka et al., Aromatic amino acid decarboxylase is involved in volatile phenylacetaldehyde production in loquat (*Eriobotrya japonica*) flowers. Plant Biotechnol. 34, 193 (2017).

78. E. Sandmeier, T. I. Hale, P. Christen, Multiple evolutionary origin of pyridoxal-5′-phosphate-dependent amino acid decarboxylases. Eur. J. Biochem. 221, 997 (1994).

