## Supplemental Material for "Ecological context determines whether inherited stress responses of hosts benefit or harm herbivores"

Alexandra Chávez *et. al*.

**This file includes:**

Figs. S1 to S6
Tables S1 to S4
References (3, 63 to 78)

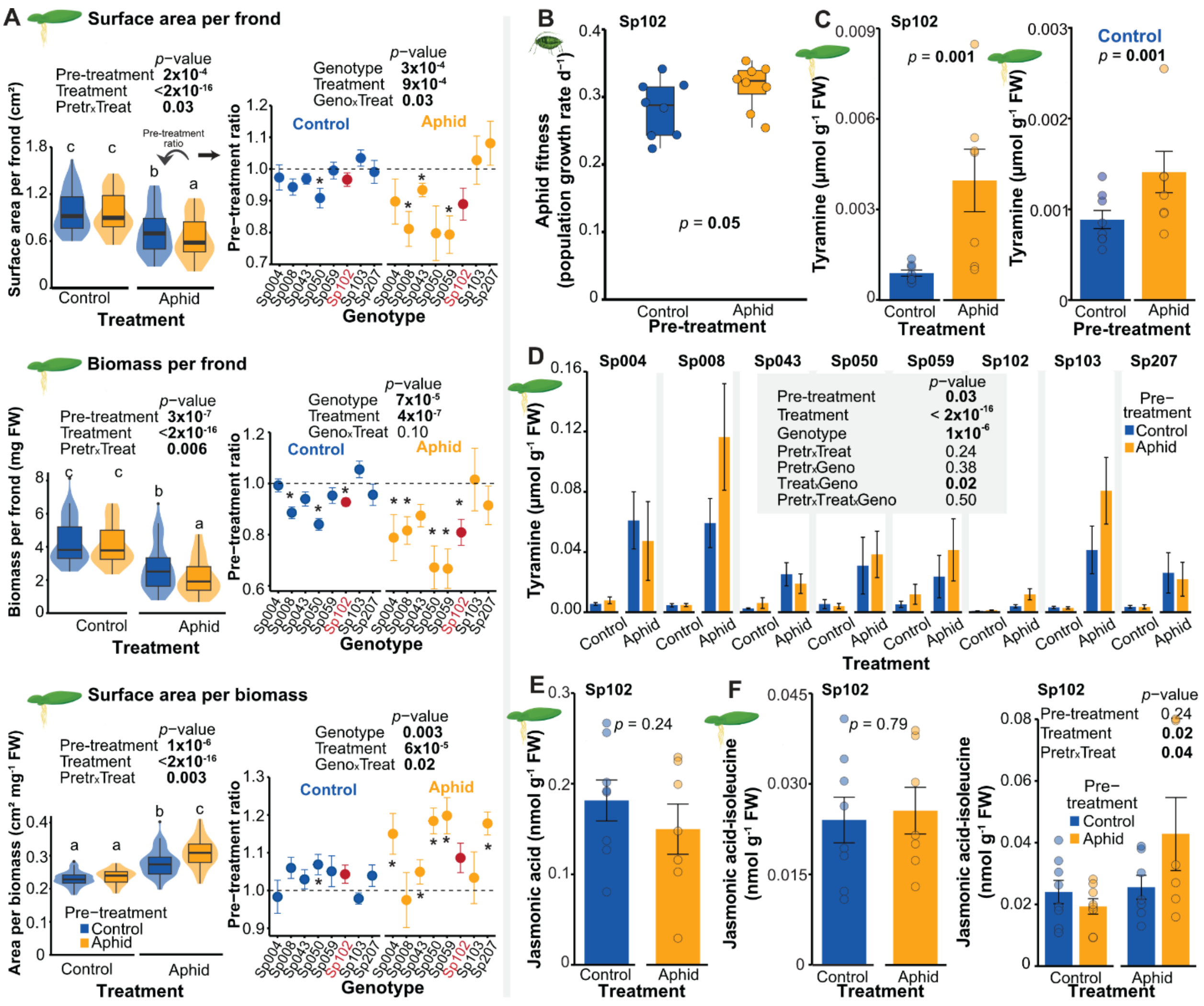

**Fig. S1.**

**Indoors, ancestral aphid herbivory by *Rhopalosiphum nymphaeae* alters the morphology and physiology of *Spirodela polyrhiza*. (A)** Ancestral aphid herbivory altered plant morphology in an environment- and genotype-dependent manner by decreasing surface area per frond and biomass per frond but increasing surface area per biomass. Morphology traits were measured after eight days of free propagation within the fitness and phenotype assay. N = 58-64 (N ~ 8 per genotype). Left section: *p*-values refer to mixed-effects models, and letters on top of violin plots indicate significant grouping by the least squares means of a post-hoc test. Right section: Pre-treatment ratios refer to the morphology of aphid pre-treated plants relative to the mean morphology of control pre-treated plants. Control and Aphid headers refer to the treatment environment within fitness and phenotype assays; error bars denote ± standard errors; the dashed line at ratio equal to one marks neutrality and asterisks denote significant deviation from neutrality (Wilcoxon rank-sum test, $P < 0.05$). *P*-values refer to mixed-effects models. (**B**) The aphid *R. nymphaeae* genotype Rn001 benefitted from previous herbivory stress on *S. polyrhiza* genotype Sp102. Aphid fitness was measured after eight days of free reproduction within the fitness and phenotype assay. *P*-values refer to a mixed-effects model. N = 8. (**C**) Aphid herbivory, as well as ancestral aphid herbivory, enhanced the concentration of tyramine in *S.*

*polyrhiza* genotype Sp102. Tyramine was measured in shoot of plants after eight days of free growth in the fitness and phenotype assay. Error bars denote ± standard errors and *p*-values refer to mixed-effects models. N = 8. Right panel: Control header refers to changes within control treatment environment within fitness and phenotype assays. (**D**) Tyramine levels in plant tissue depended on the plant genotype, the treatment environment, as well as the pre-treatment. Tyramine was measured in shoot of plants after eight days of free growth in the fitness and phenotype assay. N = 8 per genotype per treatment. (**E**) Aphid herbivory did not alter the concentrations of jasmonic acid. Jasmonic acid was measured in shoot of plants after eight days of free growth in the fitness and phenotype assay. Error bars denote ± standard errors and *p*-values refer to mixed-effects models. N = 8. (**F**) Whereas jasmonic acid-isoleucine was not induced under aphid herbivory, ancestral herbivory transgenerationally primed its concentrations in genotype Sp102 under recurrent herbivory. Jasmonic acid-isoleucine was measured in shoot of plants after eight days of free growth in the fitness and phenotype assay. Error bars denote ± standard errors and *p*-values refer to mixed-effects models. N = 8. Bold values highlight significant *p*-values. Geno = genotype, Pretreat = pre-treatment, Treat = treatment, FW = fresh weight.

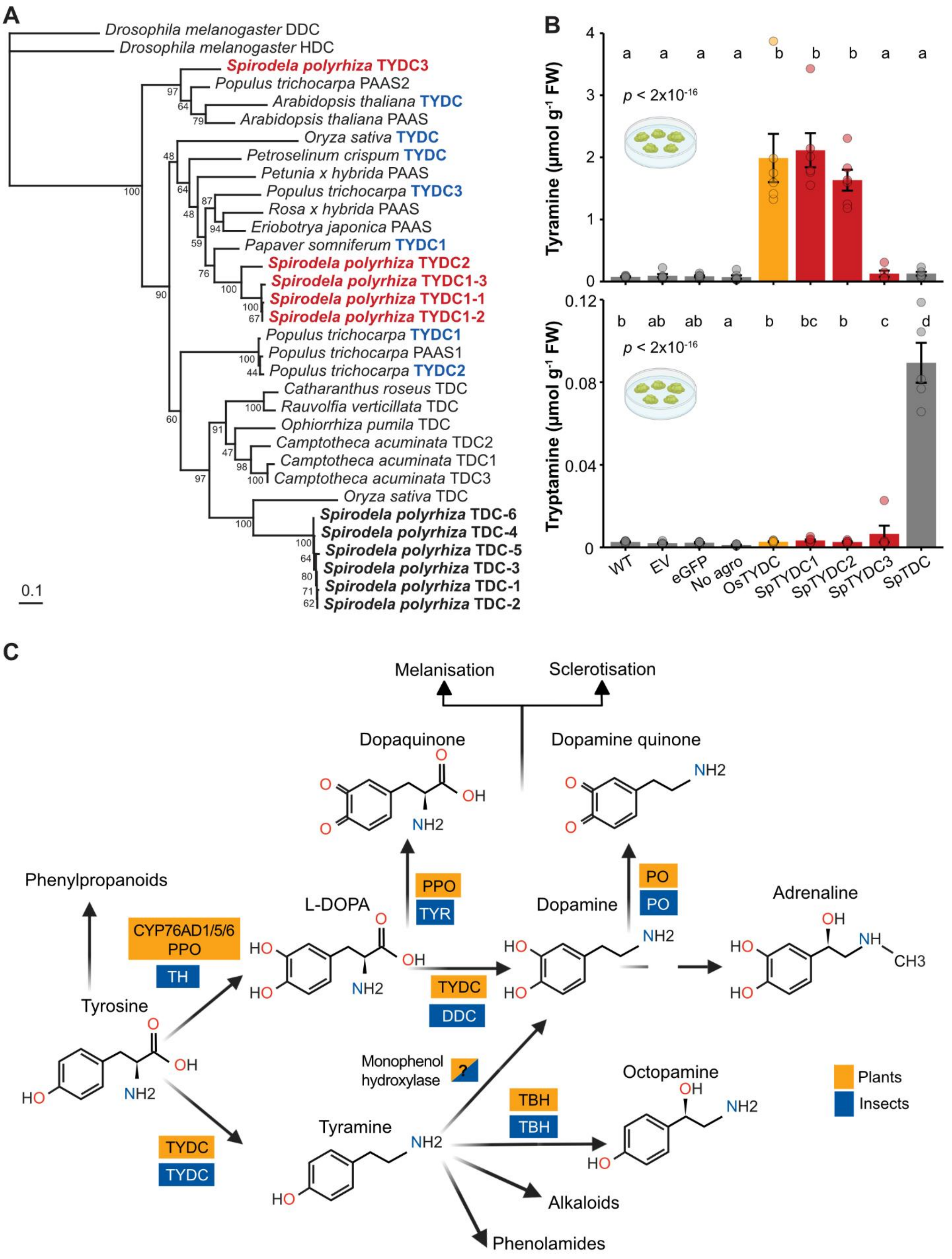

A
Drosophila melanogaster DDC
Drosophila melanogaster HDC
Spirodela polyrhiza TYDC3
Populus trichocarpa PAAS2
Arabidopsis thaliana TYDC
Arabidopsis thaliana PAAS
Oryza sativa TYDC
Petroselinum crispum TYDC
Petunia x hybrida PAAS
Populus trichocarpa TYDC3
Rosa x hybrida PAAS
Eriobotrya japonica PAAS
Papaver somniferum TYDC1
Spirodela polyrhiza TYDC2
Spirodela polyrhiza TYDC1-3
Spirodela polyrhiza TYDC1-1
Spirodela polyrhiza TYDC1-2
Populus trichocarpa TYDC1
Populus trichocarpa PAAS1
Populus trichocarpa TYDC2
Catharanthus roseus TDC
Rauvolfia verticillata TDC
Ophiorrhiza pumila TDC
Camptotheca acuminata TDC2
Camptotheca acuminata TDC1
Camptotheca acuminata TDC3
Oryza sativa TDC
Spirodela polyrhiza TDC-6
Spirodela polyrhiza TDC-4
Spirodela polyrhiza TDC-5
Spirodela polyrhiza TDC-3
Spirodela polyrhiza TDC-1
Spirodela polyrhiza TDC-2
0.1
B
Tyramine (µmol g-1 FW)
p < 2x10-16
a a a a b b b a a
Tryptamine (µmol g-1 FW)
b ab ab a b bc b c d
WT
EV
eGFP
No agro
OsTYDC
SpTYDC1
SpTYDC2
SpTYDC3
SpTDC
C
Melanisation
Sclerotisation
Dopaquinone
Dopamine quinone
Phenylpropanoids
L-DOPA
Dopamine
Adrenaline
CYP76AD1/5/6
PPO
TH
PPO
TYR
PO
PO
TYDC
DDC
Tyrosine
Monophenol
hydroxylase
?
Octopamine
TBH
TBH
TYDC
TYDC
Tyramine
Alkaloids
Phenolamides
Plants
Insects

**Fig. S2.**

**Tyrosine metabolism in plants and insects. (A)** Phylogenetic tree of characterized aromatic amino acid decarboxylases (Table S3) clustered S. polyrhiza proteins annotated as L-tyrosine decarboxylases with other L-tyrosine decarboxylases and phenylacetaldehyde synthases. L-tyrosine decarboxylases are written in blue; the three annotated *S. polyrhiza* L-tyrosine decarboxylases and their tandem copies are shown in bold red, while the *S. polyrhiza* L-tryptophan decarboxylase and the tandem copies are in bold black. (**B**) Two of the three *S. polyrhiza* L-tyrosine decarboxylase candidates, *SpTYDC1* and *SpTYDC2*, encode the synthesis of tyramine. SpTDC decarboxylates tryptophan into tryptamine. Coding sequences were transiently expressed in *S. polyrhiza* calli. A tyrosine decarboxylase from rice (*OsTYDC*) (Park, Lee, Kim, Chi, Shin and Back (*50*)) served as a positive control. (**C**) Selected aspects of tyrosine metabolism in plants and insects. Enzymes synthetizing each metabolite are written on yellow and blue rectangles. Enzyme names were taken from: Xu, Fang, Li, Yang and Chen (*60*), Brandau and Axelrod (*61*), Soares, Rogério, Cássia, Rogério, Wanderley and and Ferrarese-Filho (*62*). AADC = aromatic L-amino acid decarboxylase, agro = co-cultivation with Agrobacterium, DDC = L-dopa decarboxylase (dopa decarboxylase), eGFP = enhanced green fluorescent protein, EV = empty vector, HDC = histidine decarboxylase, Os = Oryza sativa, PAAS = phenylacetaldehyde synthase, PO = phenoloxidase or dopamine quinone synthase, PPO = polyphenol oxidase, TBH = tyramine β-hydroxylase, TDC = tryptophan decarboxylase, TH = tyrosine hydroxylase, TYDC = tyrosine decarboxylase, TYR = tyrosinase, WT = wild type.

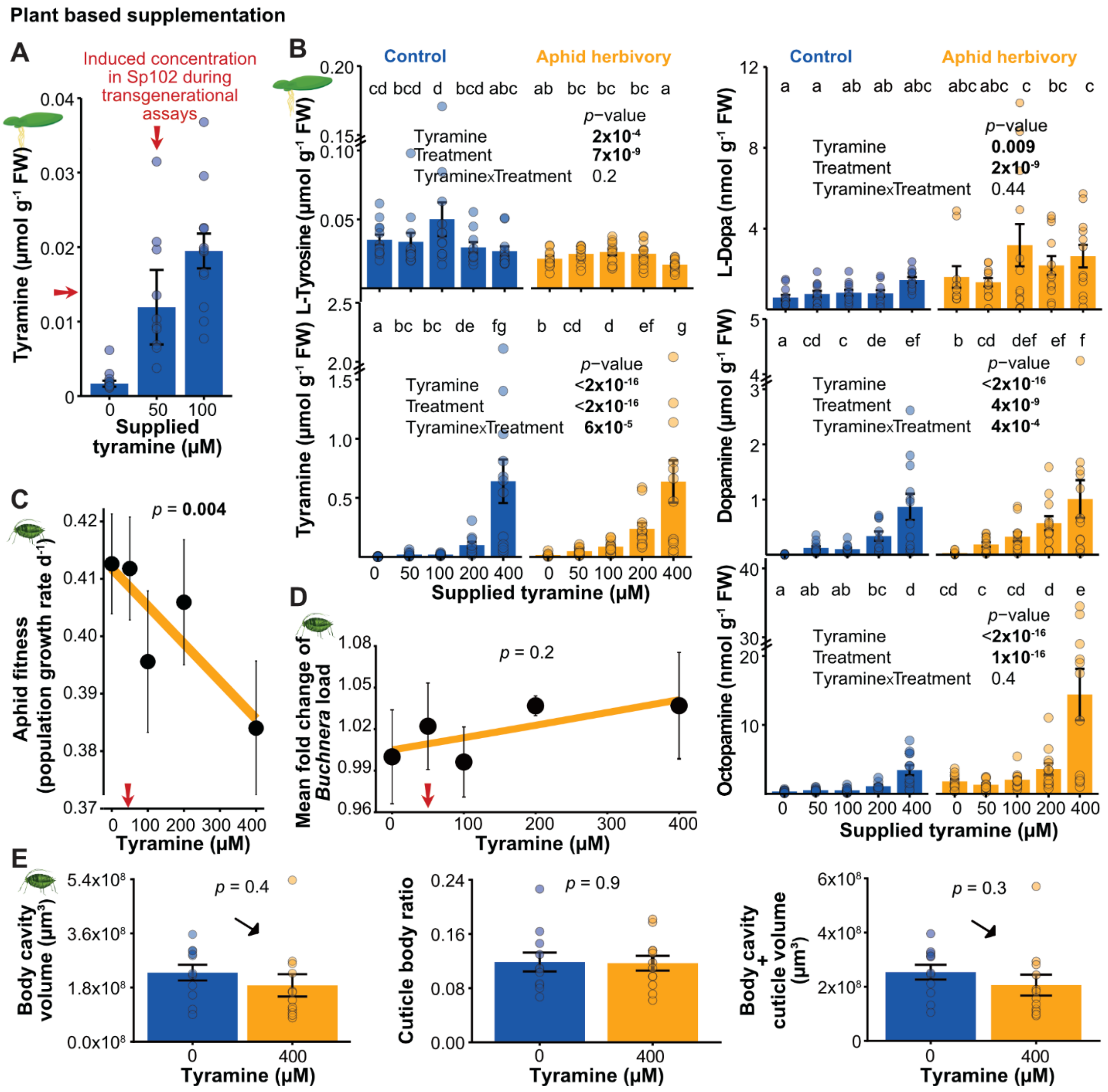


**Fig. S3.**

**Tyramine supplementation in *Spirodela polyrhiza* increases *in planta* levels of tyramine and related metabolites but does not benefit aphids.** (**A**) Supplementing *S. polyrhiza* with tyramine by adding 50-100 µM to the medium enhanced tyramine concentrations in plant shoots, reaching similar levels to those observed in transgenerational experiments (red arrow). (**B**) Supplementation with tyramine increased dopamine and octopamine accumulation in plant shoots, but these changes were not reflected in L-tyrosine, and only marginally in L-Dopa. Units for L-Dopa, dopamine and octopamine are estimated based on phenylalanine concentrations, but do not represent absolute values. *P*-values refer to mixed-effects models and letters refer to least squares means of post hoc tests. N = 12. (**C**) Supplementation of tyramine in a sucrose solution decreased the number of offspring per aphid after seven days of feeding. Each dot refers to the mean value per concentration. *P*-value refers to a linear regression. N = 5 per tyramine concentration. (**D**) The amount of *Buchnera spp*. load did not change across tyramine concentrations. Y axis represents the mean fold change (Delta Ct) of *Buchnera spp*. quantified through 16S rRNA relative to the housekeeping gene Ef1α (*55*). *P*-value refers to a linear regression. N = 4 per tyramine concentration. (**E**) Supplementation of tyramine to *S. polyrhiza* did not alter aphid cuticle traits. Arrows show the tendency of body size decrease under 400 µM in comparison to 0 µM tyramine supplementation. *P*-values refer to linear models. N = 11-12. FW = fresh weight.

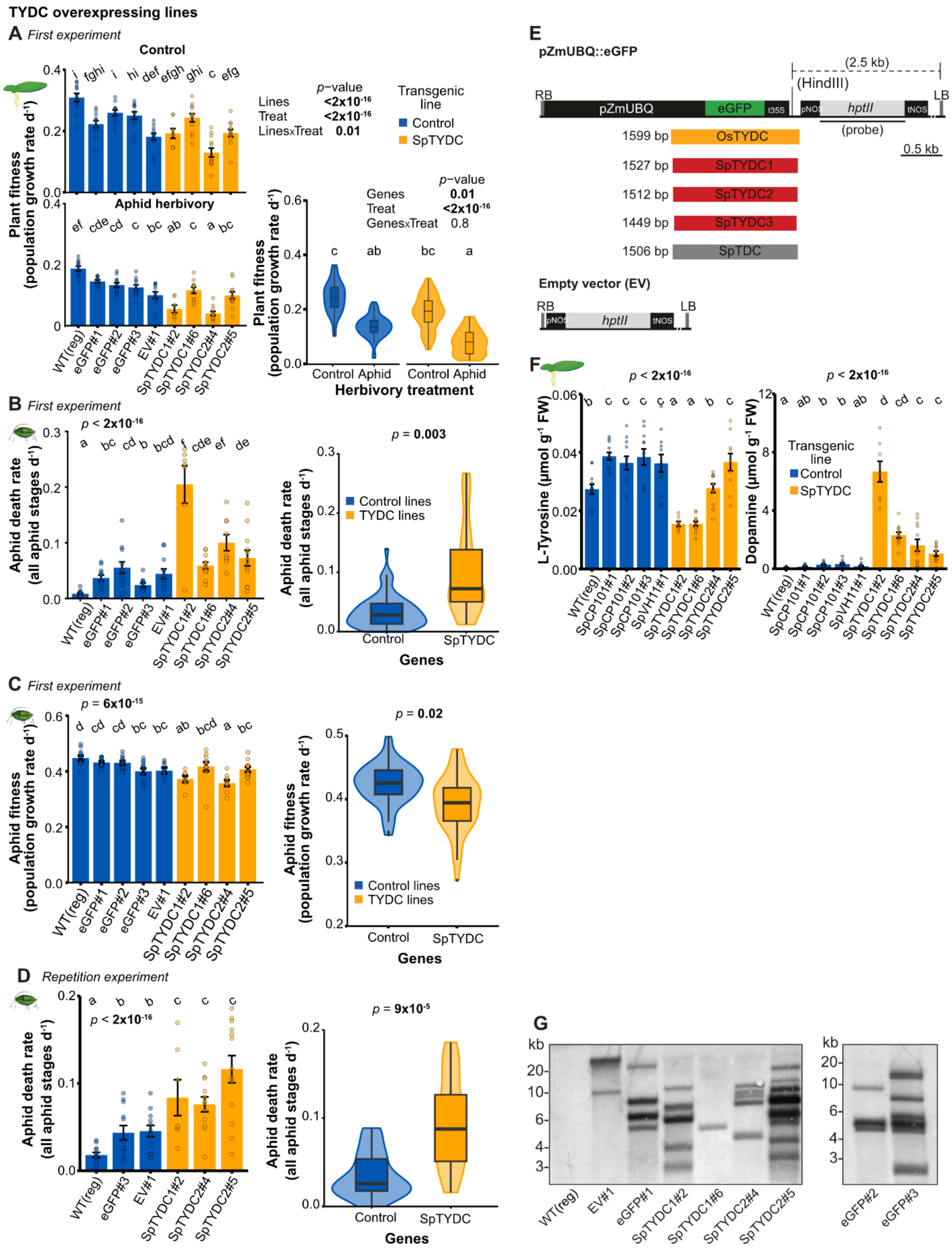
TYDC overexpressing lines
A First experiment
Control
Aphid herbivory
Plant fitness (population growth rate d-1)
p-value
Lines <2x10-16
Treat <2x10-16
LinesxTreat 0.01
Transgenic line
Control
SpTYDC
Genes 0.01
Treat <2x10-16
GenesxTreat 0.8
Control Aphid
Herbivory treatment
WT(reg) eGFP#1 eGFP#2 eGFP#3 EV#1 SpTYDC1#2 SpTYDC1#6 SpTYDC2#4 SpTYDC2#5
B First experiment
p < 2x10-16
Aphid death rate (all aphid stages d-1)
p = 0.003
Control lines
TYDC lines
Control
SpTYDC
Genes
C First experiment
p = 6x10-15
Aphid fitness (population growth rate d-1)
p = 0.02
D Repetition experiment
p = 9x10-5
E
pZmUBQ::eGFP
(2.5 kb)
(HindIII)
RB
pZmUBQ
eGFP
t35S
pNOS
hptII
tNOS
LB
(probe)
0.5 kb
1599 bp OsTYDC
1527 bp SpTYDC1
1512 bp SpTYDC2
1449 bp SpTYDC3
1506 bp SpTDC
Empty vector (EV)
F
p < 2x10-16
L-Tyrosine (µmol g-1 FW)
Dopamine (µmol g-1 FW)
WT(reg) SpCP101#1 SpCP101#2 SpCP101#3 SpVH11#1 SpTYDC1#2 SpTYDC1#6 SpTYDC2#4 SpTYDC2#5
G
kb
20- 10- 6- 4- 3-
WT(reg) EV#1 eGFP#1 SpTYDC1#2 SpTYDC1#6 SpTYDC2#4 SpTYDC2#5
eGFP#2 eGFP#3

**Fig. S4.**

**TYDC overexpression increases endogenous tyramine levels and aphid mortality. (A)** Overaccumulation of tyramine in TYDC overexpressing lines reduced plant fitness in comparison to control transgenic plants. Plant fitness was measured as increase in surface area after seven days of free propagation within fitness and phenotype assays. *P*-values refer to mixed-effects models and letters refer to least square means of post hoc tests. N = 12. (**B**) TYDC lines increased the mortality rate in aphids *R. nymphaeae* genotype Rn001 in comparison to aphids feeding on control plants. Death rate was measured considering all aphid ages after seven days of free reproduction on the transgenic lines. *P*-values refer to mixed-effects models and letters refer to least squares means of a post hoc test. N = 12. (**C**) TYDC lines decreased fitness in aphids *R. nymphaeae* genotype Rn001 in comparison to aphids feeding on control plants. Fitness (population growth rates) was measured considering all aphid ages after seven days of free reproduction on the transgenic lines. *P*-values refer to mixed-effects models and letters refer to least squares means of a post hoc test. N = 12. (**D**) Repeating the experiment showed consistent increase in aphid death rate along the TYDC lines in comparison with the control lines. Death rate was measured considering all aphid ages after seven days of free reproduction on the transgenic lines. *P*-values refer to mixed-effects models and letters refer to least squares means of a post hoc test. N = 12. (**E**) Construct maps of the T-DNA binary vectors used for *S. polyrhiza* transformation. The vector *pZmUBQ::eGFP* was used as control and backbone to clone the genes of interest, replacing the eGFP in the plant expression cassette. An additional control vector was generated by removing the eGFP expression cassette (empty vector, EV), i.e., containing only the plant selectable marker cassette *pNOs::hptII::tNOS*, which confers resistance to the antibiotic hygromycin. (**F**) Transgenic *S. polyrhiza* Sp162 lines that stably overexpress tyrosine decarboxylases (depicted in yellow) accumulate higher levels of dopamine than controls (depicted in blue). Tyramine concentrations were measured in shoots of the transgenic Sp162 after seven days of free growth. Tyrosine decarboxylases were expressed under the control of the constitutive *Zea mays* promoter ZmUBQ10. Independent transgenic lines are indicated by hash numbers. N = 12. (**G**) Southern blot of the transgenic lines. hptII = hygromycin-B-phosphotransferase, LB = left border, Os = *Oryza sativa*, pNOS = nopaline synthase promoter, pZmUBQ = *Zea mays* ubiquitin promoter, RB = right border, Sp = *Spirodela polyrhiza*, t35S = cauliflower mosaic virus 35S terminator, TDC = tryptophan decarboxylase, tNOS = nopaline synthase terminator, TYDC = tyrosine decarboxylase, FW = fresh weight.

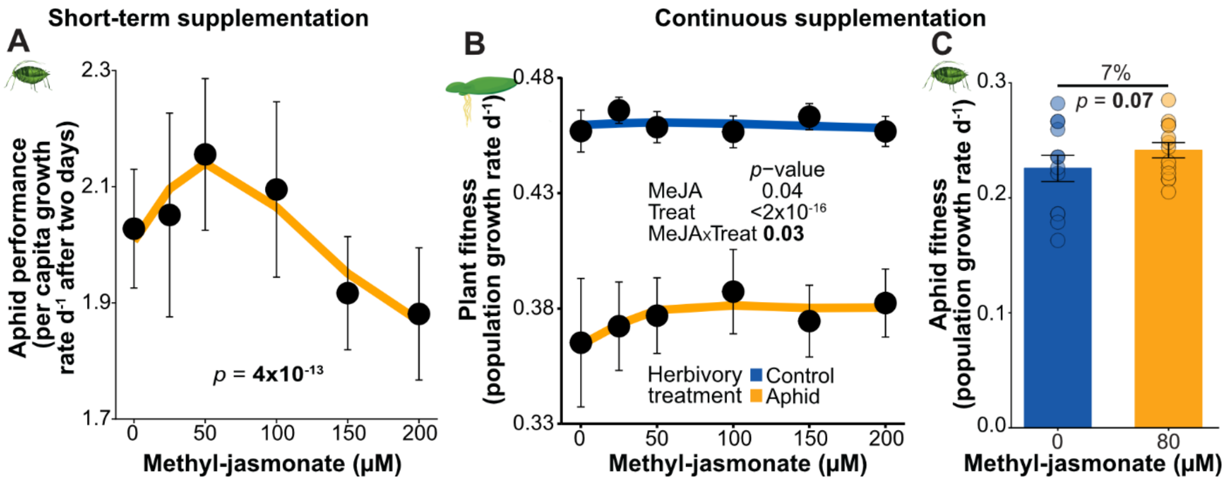


**Fig. S5.**

**Aphids benefit from moderate, but not high, levels of jasmonate induction.** (**A**) Aphid performance (per capita growth rates) increased under mild methyl-jasmonate concentrations (~50 µM) but was suppressed under higher concentrations. Performance of *R. nymphaeae* Rn001 was measured over 48 h in the absence of methyl-jasmonate following three days of induction with methyl-jasmonate in *S. polyrhiza* Sp102. Dots represent mean values and error bars denote ± standard errors; *p*-value at the bottom refers to a mixed-effects model with natural spline. N = 16 per concentration. (**B**) High methyl-jasmonate concentrations protected plants under aphid herbivory. Plant fitness was measured as increased in surface area across eight days of continuous methyl-jasmonate induction. Error bars denote ± standard errors; *p*-value refers to a mixed-effects model. N = 16. (**C**) Mild methyl-jasmonate doses tended to increase aphid fitness under continuous methyl-jasmonate supplementation. Aphid fitness (population growth rates) was measured over eight days of continuous methyl-jasmonate induction in Sp102 plants. Error bars denote ± standard errors; *p*-value refers to a mixed-effects model. N = 12.

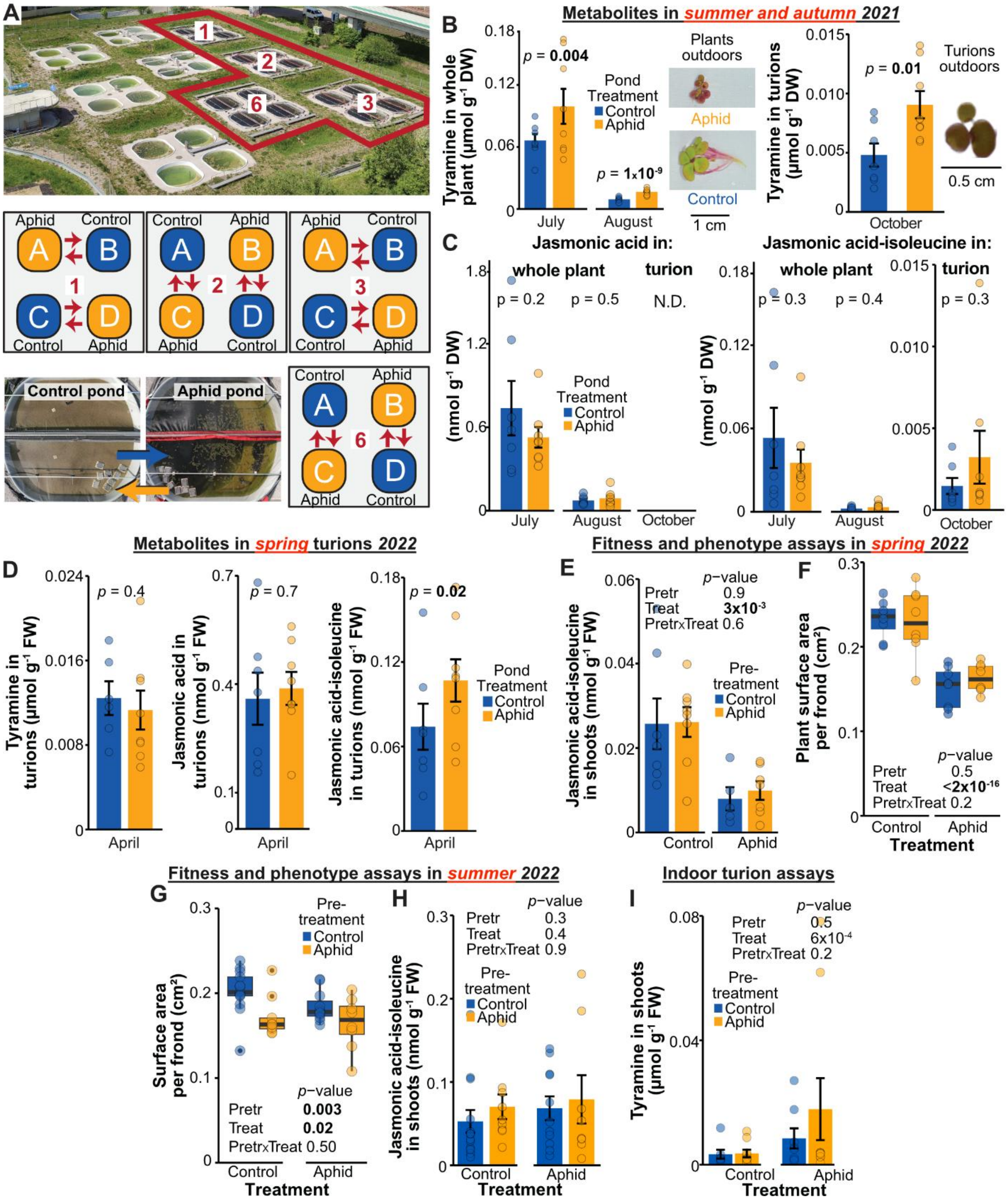


**Fig. S6.**

**Outdoors, transgenerational plasticity in tyramine levels and plant phenotypes—but not jasmonic acid—persists within, but not across, growing seasons. (A**) Spatial arrangement of control and herbivory ponds in which populations of the monoclonal *S. polyrhiza* genotype Sp102 were grown across two growing seasons with contrasting levels of herbivory by the aphid *R. nymphaeae* genotype Rn001. Red lines delimit the experimental ponds. Arrows indicate the direction in which plants were moved between pairs of ponds for the transplant experiments. Overview of two ponds shows how the transplanted plants were cultivated within the reciprocal pond. (**B**) Concentrations of tyramine increased under aphid herbivory during the first growing season of the outdoor experiments, being retained in the turions. Tyramine was measured in the whole plant in July 2021 and in turions in August 2021. Aphid herbivory reduced frond and root size and increased red coloration in shoots of the population. *P*-values refer to mixed-effects models. N = 8 per treatment. (**C**) Jasmonates were not affected by aphid herbivory during the first growing season of the outdoor experiment. Jasmonates were measured in whole plant and turions in July and August 2021, respectively. *P*-values refer to mixed-effects models. N = 8 per treatment. (**D**) Physiological changes induced by aphid herbivory did not persist in turions that emerged the following season. Metabolites were measured in floating turions before germination in April 2022. Error bars refer to ± standard errors; *p*-values refer to mixed-effects models. N = 8 per treatment. (**E**) Herbivory-induced changes in jasmonic acid-isoleucine within turions were not transmitted to the plants. *S. polyrhiza* genotype Sp102 was cultivated for three weeks under control conditions and subsequently allowed to grow freely for an additional three weeks in the presence or absence of herbivory, after which metabolites were quantified. Assays were performed in spring 2022. Error bars refer to ± standard errors; *p*-values refer to mixed-effects models. N = 8 independent replicates. (**F**) Herbivory in the previous growing season did not alter surface area per frond in plants. *S. polyrhiza* genotype Sp102 was cultivated for three weeks under control conditions and subsequently allowed to grow freely for an additional three weeks under aphid herbivory or control environment, environments in which fitness and morphology were measured. Assays were performed in spring 2022. *P*-values refer to mixed-effects models. N = 8 independent replicates (**G**) Ancestral aphid herbivory within growing season transgenerationally decreased surface area per frond on *S. polyrhiza* genotype Sp102 regardless of the environment. Plants grew freely for three weeks under control conditions, previous to fitness and phenotype assays under control conditions and aphid herbivory. Assays were performed in summer 2022. Error bars refer to ± standard errors; *p*-values refer to a mixed-effects model. N = 8 independent replicates. (**H**) Ancestral aphid herbivory did not alter jasmonic acid-isoleucine concentrations. Jasmonic acid-isoleucine was measured in shoot material of plants that reproduced freely for three weeks under control and aphid herbivory after three weeks of recovery from stress. Error bars refer to ± standard errors; *p*-values refer to mixed-effects models. N = 8 independent replicates per pond treatment. (**I**) Indoors, transgenerationally primed tyramine concentrations did not persist after turion formation. Turion formation was induced in Sp102 three generations after aphid herbivory pre-treatment, and plants germinating from the turions were used to continue the transgenerational experiments. Tyramine was measured in shoot of plants. Error bars refer to ± standard errors; *p*-values refer to a mixed-effects model.  N = 8. DW = dry weight, FW = fresh weight, N.D. = no detected.

### Table S1.

### Description of the L-tyrosine and L-tryptophan decarboxylase genes in *Spirodela polyrhiza*.

| Gene | Copy | Chromosome | N° exons | Start position | Ending position | Gene strand | Exon length | Accession 1 | Accession 2 |
|---|---|---|---|---|---|---|---|---|---|
| **SpTYDC1** | SpTYDC1_1 | 15 | 1 | 2759464 | 2760990 | + | 1530 | CM046543.1 | JANEYH010000015.1 |
| | *SpTYDC1p_1* | | 0 | *2763644* | *2763961* | + | *(858-1167)* | | |
| | *SpTYDC1p_2* | | 0 | *2764475* | *2764791* | + | *(1197-1515)* | | |
| | SpTYDC1_2 | | 1 | 2768337 | 2769863 | + | 1530 | | |
| | SpTYDC1_3 | | 1 | 2774914 | 2776443 | + | 1530 | | |
| **SpTYDC2** | SpTYDC2 | 3 | 1 | 2758987 | 2760498 | + | 1512 | CM046531.1 | JANEYH010000003.1 |
| | *SpTYDC2p1* | | 0 | *2766571* | *2767158* | + | *(588)* | | |
| **SpTYDC3** | SpTYDC3 | 9 | 13 | 329615 | 329716 | + | 1-102 | CM046537.1 | JANEYH010000009.1 |
| | | | | 330087 | 330159 | + | 103-175 | | |
| | | | | 330316 | 330546 | + | 176-406 | | |
| | | | | 331510 | 331673 | + | 407-570 | | |
| | | | | 332102 | 332242 | + | 571-711 | | |
| | | | | 332898 | 332957 | + | 712-771 | | |
| | | | | 334305 | 334469 | + | 772-936 | | |
| | | | | 335088 | 335144 | + | 937-993 | | |
| | | | | 335568 | 335632 | + | 994-1058 | | |
| | | | | 335876 | 335999 | + | 1059-1182 | | |
| | | | | 336120 | 336257 | + | 1183-1320 | | |
| | | | | 336336 | 336464 | + | 1321-1449 | | |
| **SpTDC** | SpTDC_1 | 3 | 2 | 8857223 | 8856138 | - | 1-1086 | CM046531.1 | JANEYH010000003.1 |
| | | | | 8855991 | 8855566 | - | 1087-1506 | | |
| | SpTDC_2 | | 2 | 8863375 | 8862290 | - | 1-1086 | | |
| | | | | 8862143 | 8861718 | - | 1087-1506 | | |
| | SpTDC_3 | | 2 | 8869415 | 8868330 | - | 1-1086 | | |
| | | | | 8868183 | 8867758 | - | 1087-1506 | | |
| | SpTDC_4 | | 2 | 8874099 | 8875184 | + | 1-1086 | | |
| | | | | 8875323 | 8875742 | + | 1087-1506 | | |
| | SpTDC_5 | | 2 | 8880807 | 8881892 | + | 1-1086 | | |
| | | | | 8882045 | 8882464 | + | 1087-1506 | | |
| | SpTDC_6 | | 2 | 8887967 | 8889052 | + | 1-1086 | | |
| | | | | 8889205 | 8889614 | + | 1087-1496 | | |

**Table S2.**

**Settings for compounds added to the LC-MS method described previously (*3*).**

| Analyte | RT [min] | Q1 [m/z] | Q3 [m/z] | Dwell time [ms] | CE [V] | Q1/Q3 Pre Bias [V] | ISTD |
|---|---|---|---|---|---|---|---|
| Kynurenic acid | 3.50 | (+) 190.05 | 144.10 | 20 | -20 | -24 / -22 | $^{13}C_9$, $^{15}N_1$-L-Tyrosine |
| | | (+) 190.05 | 89.05 | 15 | -40 | -23 / -16 | |
| | | (+) 190.05 | 116.15 | 15 | -32 | -13 / -11 | |
| Kynurenine | 2.515 | (+) 209.09 | 192.10 | 20 | -10 | -10 / -20 | $^{13}C_9$, $^{15}N_1$-L-Tyrosine |
| | | (+) 209.09 | 146.10 | 15 | -19 | -10 / -15 | |
| Serotonin | 2.120 | (+) 177.10 | 160.10 | 20 | -14 | -30 / -30 | $^{13}C_9$, $^{15}N_1$-L-Tyrosine |
| | | (+) 177.10 | 115.10 | 15 | -28 | -22 / -23 | |
| Dopamine | 0.990 | (+) 154.09 | 91.10 | 20 | -24 | -11 / -19 | $^{13}C_9$, $^{15}N_1$-L-Tyrosine |
| | | (+) 154.09 | 137.20 | 15 | -15 | -11 / -14 | |
| L-Dopa | 0.930 | (+) 198.08 | 135.15 | 15 | -20 | -14 / -25 | $^{13}C_9$, $^{15}N_1$-L-Tyrosine |
| | | (+) 198.08 | 152.00 | 20 | -14 | -13 / -16 | |
| Norepinephrine | 0.570 | (+) 170.08 | 107.10 | 20 | -21 | -12 / -11 | $^{13}C_9$. $^{15}N_1$-L-Tyrosine |
| | | (+) 170.08 | 77.10 | 15 | -37 | -12 / -14 | |
| Epinephrine | 0.615 | (+) 184.10 | 166.15 | 20 | -12 | -14 / -22 | $^{13}C_9$. $^{15}N_1$-L-Tyrosine |
| | | (+) 184.10 | 107.10 | 15 | -21 | -14 / -21 | |
| Octopamine | 0.610 | (+) 154.09 | 91.15 | 20 | -22 | -11 / -18 | $^{13}C_9$. $^{15}N_1$-L-Tyrosine |
| | | (+) 154.09 | 119.15 | 15 | -18 | -12 / -25 | |
| Melatonin | 3.880 | (+) 233.13 | 174.10 | 20 | -15 | -20 / -20 | $^{13}C_9$. $^{15}N_1$-L-Tyrosine |
| | | (+) 233.13 | 130.15 | 15 | -43 | -11 / -23 | |
| | | (+) 233.13 | 159.20 | 15 | -26 | -11 / -30 | |
| GABA | 0.535 | (+) 104.07 | 87.10 | 20 | -13 | -10 / -16 | $^{13}C_9$. $^{15}N_1$-L-Tyrosine |
| | | (+) 104.07 | 69.10 | 15 | -16 | -22 / -12 | |

RT: retention time.
CE: collision energy.
ISTD: internal standard
Qualifiers are highlighted in grey.

### Table S3.
### Genes used in the phylogenetic analysis of the aromatic decarboxylases.

| | | | | | | Decarboxylation | | | | **Decarboxylation + Transamination** | | | |
|---|---|---|---|---|---|---|---|---|---|---|---|---|---|
| **Organism** | **Abbreviation based on activity** | **Abbreviation in reference** | **Genbank accession (Gene ID)** | **Assay** | **Reference** | tyrosine → tyramine | Tryptophan → tryptamine | phenyl alanine → phenyl-ethylamine | L-Dopa → dopamine | phenyl alanine → phenyl-acetaldehyde | Tyrosine → 4-OH-phenyl-acetaldehyde | Tryptophan → indole-3-acetaldehyde | L-Dopa → L-Dopaldehyde |
| *Arabidopsis thaliana* | TYDC | | AEE85523 (AT4G28680) | purified recombinant $His_6$-protein | Lehmann and Pollmann (*63*) | **yes** | no | no | no | | | | |
| *Oryza sativa* | TYDC | | AK065830 / BAG89694 | tyramine in overexpressing rice leaves | Lee, Kang, Park, Park and Back (*64*). Park, Lee, Kim, Chi, Shin and Back (*50*). Facchini and De Luca (*65*) | **yes** | | | | | | | |
| *Papaver somniferum* | TYDC1 | | AAC61844 | recombinant protein *E. coli* extract | Facchini and De Luca (*66*) | **yes** | no | very low | **yes** | | | | |
| *Petroselinum crispum* | TYDC | | AAA33860 | recombinant protein *E. coli* extract | Kawalleck, Keller, Hahlbrock, Scheel and Somssich (*67*) | **yes** | no | very low | low | | | | |
| *Populus trichocarpa* | TYDC1 | AADC1 | PNT06818 (Potri.013G052800) | purified recombinant $His_6$-protein | Günther, Lackus, Schmidt, Huber, Stödler, Reichelt, Gershenzon and Köllner (*68*) | **yes** | yes | **yes** | | | | | |
| *Populus trichocarpa* | TYDC2 | AADC2 | PNS99079 (Potri.016G114300) | purified recombinant $His_6$-protein | Günther, Lackus, Schmidt, Huber, Stödler, Reichelt, Gershenzon and Köllner (*68*) | **yes** | no | low | | | | | |
| *Populus trichocarpa* | TYDC3 | AADC3 | PNT39402 (Potri.004G036200) | purified recombinant $His_6$-protein | Günther, Lackus, Schmidt, Huber, Stödler, Reichelt, Gershenzon and Köllner (*68*) | **yes** | no | yes | | | | | |
| *Camptotheca acuminata* | TDC1 | | AAB39708 | recombinant protein *E. coli* extract | López-Meyer and Nessler (*69*) | no | **yes** | no | no | | | | |
| *Camptotheca acuminata* | TDC2 | | AAB39709 | recombinant protein *E. coli* extract | López-Meyer and Nessler (*69*) | no | **yes** | no | no | | | | |
| *Camptotheca acuminata* | TDC3 | | UZH44855 | purified recombinant $His_6$-protein | Qiao, Chen, Liu, Huang, Li, Zhang and Luo (*70*) | | **yes** | | | | | | |
| *Catharanthus roseus* | TDC | | P17770.1 / TDC_CATRO | recombinant protein *E. coli* extract | De Luca, Marineau and Brisson (*71*) | | **yes** | | | | | | |
| *Ophiorrhiza pumila* | TDC | | BAC41515 | purified recombinant $His_6$-protein | Yamazaki, Sudo, Yamazaki, Aimi and Saito (*72*) | | **yes** | | | | | | |

| | | | | | | | | | | | | | |
|---|---|---|---|---|---|---|---|---|---|---|---|---|---|
| *Oryza sativa* | TDC | | BAG91223 | purified recombinant $His_6$-protein | Kang, Jeong, Lee and Natarajan (*73*) | no | **yes** | no | no | | | | |
| *Rauvolfia verticillata* | TDC | | ADL28270 | purified recombinant $His_6$-protein | Liu, Chen, Chen, Zhang, Peng, Yang, Ming, Lan and Liao (*74*) | | **yes** | | | | | | |
| *Arabidopsis thaliana* | PAAS | | Q8RY79 / PAAS_ARATH (AT2G20340) | purified recombinant $His_6$-protein | Gutensohn, Klempien, Kaminaga, Nagegowda, Negre-Zakharov, Huh, Luo, Weizbauer, Mengiste, Tholl and Dudareva (*75*) | | | | | **yes** | no | no | **yes** |
| *Petunia hybrida* | PAAS | | ABB72475 | purified recombinant $His_6$-protein | Kaminaga, Schnepp, Peel, Kish, Ben-Nissan, Weiss, Orlova, Lavie, Rhodes, Wood, Porterfield, Cooper, Schloss, Pichersky, Vainstein and Dudareva (*76*) | | | | | **yes** | no | no | no |
| *Rosa hybrida* | PAAS | | ABB04522 | purified recombinant $His_6$-protein | Kaminaga, Schnepp, Peel, Kish, Ben-Nissan, Weiss, Orlova, Lavie, Rhodes, Wood, Porterfield, Cooper, Schloss, Pichersky, Vainstein and Dudareva (*76*) | | | | | **yes** | no | no | no |
| *Eriobotrya japonica* | PAAS | AADC1 | BBE38027 | purified recombinant $His_9$-protein | Koeduka, Fujita, Furuta, Suzuki, Tsuge and Matsui (*77*) | | | | | **yes** | | | |
| *Populus trichocarpa* | PAAS1 | AAS1 | PNT06819 (Potri.013G052900) | purified recombinant $His_6$-protein | Günther, Lackus, Schmidt, Huber, Stödler, Reichelt, Gershenzon and Köllner (*68*) | | | | | **yes** | no | no | |
| *Populus trichocarpa* | PAAS2 | AAS2 | KAI9400498 (Potri.002G255600) | purified recombinant $His_6$-protein | Günther, Lackus, Schmidt, Huber, Stödler, Reichelt, Gershenzon and Köllner (*68*) | | | | | **yes** | yes | yes | |
| *Drosophila melanogaster* | DDC | | P05031 | DDC = dopa decarboxylase | Sandmeier, Hale and Christen (*78*) | | | | | | | | |
| *Drosophila melanogaster* | HDC | | X70644 (CAA49989) | HDC = histidine decarboxylase | Sandmeier, Hale and Christen (*78*) | | | | | | | | |

**Table S4.**

**Summary of the RNAseq data quality.**

| Treatment | Library | Raw reads | Raw bases | Clean reads | Clean bases | Error rate | $Q_{Phred}$ 20 | $Q_{Phred}$ 30 | GC_pct |
|---|---|---|---|---|---|---|---|---|---|
| Control | CtrCtr102_1 | 36439634 | 5.47G | 35332794 | 5.3G | 0.02 | 97.01 | 91.52 | 57.57 |
| Recurrent control | AphCtr102_1 | 31717432 | 4.76G | 30867196 | 4.63G | 0.02 | 96.79 | 91.01 | 58.47 |
| First time herbivory | CtrAph102_1 | 30637166 | 4.6G | 29639046 | 4.45G | 0.01 | 96.97 | 91.36 | 57.31 |
| Recurrent herbivory | AphAph102_1 | 28618342 | 4.29G | 27456896 | 4.12G | 0.02 | 96.28 | 89.92 | 57.35 |
| Control | CtrCtr102_2 | 25575982 | 3.84G | 24612236 | 3.69G | 0.02 | 95.91 | 88.99 | 58.82 |
| Recurrent control | AphCtr102_2 | 65998370 | 9.9G | 62854542 | 9.43G | 0.02 | 96.91 | 91.31 | 57.76 |
| First time herbivory | CtrAph102_2 | 44029604 | 6.6G | 42909840 | 6.44G | 0.02 | 96.62 | 90.15 | 57.54 |
| Recurrent herbivory | AphAph102_2 | 39795074 | 5.97G | 38445924 | 5.77G | 0.02 | 96.79 | 90.94 | 57.62 |
| Control | CtrCtr102_4 | 46814044 | 7.02G | 45319562 | 6.8G | 0.02 | 96.91 | 91.33 | 58.38 |
| Recurrent control | AphCtr102_4 | 42940640 | 6.44G | 41608930 | 6.24G | 0.02 | 96.88 | 91.24 | 57.92 |
| First time herbivory | CtrAph102_4 | 33545628 | 5.03G | 32429206 | 4.86G | 0.01 | 97.12 | 91.58 | 57.11 |
| Recurrent herbivory | AphAph102_4 | 34608740 | 5.19G | 33403204 | 5.01G | 0.02 | 96.7 | 90.76 | 57.97 |
| Control | CtrCtr102_5 | 36237892 | 5.44G | 34761616 | 5.21G | 0.02 | 96.35 | 89.82 | 58.05 |
| Recurrent control | AphCtr102_5 | 43301092 | 6.5G | 42014790 | 6.3G | 0.02 | 96.81 | 90.98 | 58.12 |
| First time herbivory | CtrAph102_5 | 32848532 | 4.93G | 31700254 | 4.76G | 0.02 | 96.68 | 90.52 | 58.17 |
| Recurrent herbivory | AphAph102_5 | 36127624 | 5.42G | 34540868 | 5.18G | 0.01 | 98.11 | 94.73 | 57.78 |
| Control | CtrCtr102_7 | 41755790 | 6.26G | 40274198 | 6.04G | 0.02 | 96.78 | 91.02 | 58.36 |
| Recurrent control | AphCtr102_7 | 45568450 | 6.84G | 43675358 | 6.55G | 0.02 | 96.91 | 91.27 | 57.94 |
| First time herbivory | CtrAph102_7 | 42187840 | 6.33G | 40566184 | 6.08G | 0.02 | 96.61 | 90.67 | 58.5 |
| Recurrent herbivory | AphAph102_7 | 41362274 | 6.2G | 39576098 | 5.94G | 0.02 | 96.71 | 90.71 | 58.29 |

**Raw reads**: Reads count from the raw data. four rows as a unit. with statistics of reads count for every sequencing.
**Raw bases**: Base number of raw data = number of raw reads * sequence length. converting unit to G.
**Clean reads**: Base number of raw data after filtering = number of clean reads * sequence length. converting unit to G.
**Clean bases**: (clean base=clean reads*150bp) number multiply read length. saved in G unit.
**Error rate**: Average sequencing error rate. which is calculated by Qphred=-10log10(e).
**$Q_{Phred}$ 20**: The percentage of the bases whose $Q_{Phred}$ values is greater than 20. (Number of bases with $Q_{Phred}$ value > 20) / (Number of total bases) *100.
**$Q_{Phred}$ 30**: The percentage of the bases whose $Q_{Phred}$ values is greater than 30. (Number of bases with $Q_{Phred}$ value > 30) / (Number of total bases) *100.
**GC pct**: The percentage of G&C base numbers of total bases.(G&C base number) / (Total base number)*10
